# Focal astrocyte Kir4.1 loss drives seizures, spreading depolarizations and postictal impairments

**DOI:** 10.64898/2026.06.15.732359

**Authors:** Neela K. Codadu, Yunan Gao, Olga Tyurikova, Zixi Dai, Xueting Ban, Yuyan Weng, Eduard Masvidal-Codina, Jose A Garrido, Anton Guimera-Brunet, Dmitri A. Rusakov, Kevan Hashemi, Nicholas D. Mazarakis, Rob C. Wykes

## Abstract

Astrocytic dysfunction is increasingly recognised as an important contributor to epileptogenesis and seizure dynamics. Kir4.1 potassium channels expressed in astrocytes play a critical role in activity-dependent extracellular potassium buffering, and their loss may impair network stability and promote pathological hyperexcitability. In epilepsy, seizures can be accompanied by spreading depolarizations (SDs), propagating waves of neuronal and glial depolarization. While seizure-associated SDs have been proposed to terminate seizures and limit seizure spread, they have also been implicated in postictal dysfunction and sudden unexpected death in epilepsy (SUDEP), and their significance in chronic epilepsy remains unclear.

Here, we tested whether focal loss of astrocytic Kir4.1 in the adult hippocampus is sufficient to disrupt potassium buffering, generate spontaneous seizures, and promote seizure-associated SDs. Using astrocyte-targeted viral vector Cre recombinase in adult Kir4.1-floxed mice, we induced focal hippocampal reduction of Kir4.1 expression. This impaired activity-dependent potassium buffering, producing enhanced stimulation-evoked extracellular potassium accumulation and larger DC shifts in hippocampal slices. Chronic wireless EEG recordings demonstrated that focal astrocytic Kir4.1 loss was sufficient to induce spontaneous recurrent seizures and interictal epileptiform activity.

To investigate seizure-associated SDs, we combined optogenetic stimulation with graphene-based micro-transistor recordings capable of stable full-bandwidth DC electrophysiology in awake head-fixed mice. Focal hippocampal loss of astrocytic Kir4.1 markedly increased susceptibility to evoked seizures accompanied by SDs. We used chronic wireless DC-coupled video-telemetry recordings to continuously monitor seizure and SD dynamics in freely moving mice. SDs occurred frequently during generalized seizures and were first detected in cortical channels. Seizures accompanied by SDs exhibited greater spectral power, longer duration, and prolonged postictal depression compared with seizures alone. Behaviourally, seizures accompanied by SDs were linked to postictal impairment characterised by behavioural arrest and abnormal motor behaviours. These findings highlight the utility of graphene micro-transistor arrays and chronic DC-coupled telemetry for resolving infraslow (<0.1 Hz) epileptic dynamics that are largely inaccessible using conventional electrophysiological approaches. Ground-truth DC-coupled recordings enabled identification of AC-band electrographic signatures that segregated seizures with SDs from seizures alone, raising the possibility that SD-associated seizures may be retrospectively inferred from conventional AC-coupled epilepsy datasets.

We demonstrate that focal astrocytic Kir4.1 loss in the adult brain impairs potassium buffering and is sufficient to drive spontaneous seizures, supporting astrocytic potassium dysregulation as a determinant of seizure and spreading depolarization susceptibility in epilepsy. Furthermore, seizure-associated SDs are strongly linked to increased postictal impairments, supporting the concept that SDs are major determinants of pathological postictal states.

## Introduction

Epilepsy is increasingly recognised as a disorder of neuron-glia network dysfunction rather than purely neuronal hyperexcitability.^1^ Astrocytes play a central role in maintaining network stability through regulation of extracellular potassium, glutamate clearance, water homeostasis, and metabolic support at tripartite synapses.^2,3^ Among the mechanisms underlying astrocytic potassium buffering, inwardly rectifying potassium (Kir) 4.1 channels are particularly important for maintaining extracellular potassium homeostasis and the hyperpolarized resting membrane potential of astrocytes.^4,5^ Kir4.1 channels are functionally coupled to glutamate uptake mechanisms and critically regulate activity-dependent synaptic transmission and network excitability.^5,6^ Consistent with its role in potassium homeostasis, Kir4.1 dysfunction or reduced expression has been associated with neuronal hyperexcitability and epilepsy in both experimental models and human epileptic brain tissue.^2,7-9^ In parallel, mutations in *KCNJ10*, the gene encoding Kir4.1, cause severe epilepsy syndromes including EAST/SeSAME syndrome, further supporting a causal relationship between Kir4.1 dysfunction and seizure susceptibility.^2,10^ Experimental studies have demonstrated that loss of Kir4.1 disrupts extracellular potassium buffering, depolarises astrocytes, impairs glutamate uptake, and promotes neuronal hyperexcitability.^4,5^ However, previous constitutive Kir4.1 knockout models exhibit severe developmental abnormalities, widespread neurological dysfunction, and early lethality, complicating interpretation of the direct contribution of acquired astrocytic Kir4.1 dysfunction to epileptogenesis in the mature brain.^4^ Consequently, whether focal acquired loss of astrocytic Kir4.1 within established adult neural circuits is sufficient to drive spontaneous seizures remains unknown.

Spreading depolarizations (SDs) are a type of slowly propagating waves of neuronal and glial depolarization accompanied by profound disturbances in ionic homeostasis and suppression of ongoing network activity.^11^ Although historically associated primarily with migraine with aura and acute brain injury, SDs are increasingly recognised in epilepsy and can occur spontaneously before, during, or following seizures in both experimental models and human epilepsy.^12-15^ Several studies have proposed that seizure-associated SDs may represent an endogenous anti-seizure mechanism capable of terminating ictal activity and transiently suppressing subsequent seizure susceptibility.^16,17^ However, SDs have also been linked to impaired postictal behavioural recovery, abnormal ambulation, and catastrophic brainstem dysfunction associated with SUDEP.^18-22^ Together, these findings underscore that SDs are neither epiphenomenal nor uniformly deleterious but rather context-dependent events whose functional significance in epilepsy remains incompletely understood. Despite growing recognition of seizure-SD coupling, the mechanisms linking astrocytic dysfunction, epileptic hyperexcitability, and SD susceptibility remain unclear.

Here, we investigated whether focal loss of astrocytic Kir4.1 within the adult hippocampus is sufficient to impair potassium buffering, generate spontaneous seizures, and promote seizure-associated SDs. Using *ex vivo* electrophysiology and potassium biosensor recordings, we first examined how astrocytic Kir4.1 loss alters activity-dependent potassium handling and neuronal excitability. We then combined optogenetic stimulations with full-bandwidth graphene-based electrophysiology^23^ to determine how Kir4.1 loss influences susceptibility to induced seizures and SDs *in vivo*. Finally, chronic DC-coupled telemetry with time-locked video monitoring in freely moving mice was used to define the natural history of spontaneous seizures and seizure-associated SDs, together with their electrophysiological and behavioural consequences.

Our findings demonstrate that astrocytic Kir4.1 loss within the adult hippocampus is sufficient to drive spontaneous recurrent seizures and promote seizure-associated SDs. SD-associated seizures were accompanied by more severe postictal electrophysiological and behavioural impairment, linking astrocytic potassium buffering dysfunction to seizure-SD coupling and impaired postictal recovery.

## Materials and methods

### Ethical approval

All animal handling and experiments were conducted according to the guidelines laid by the UK Home Office and Animals (Scientific Procedures) Act 1986, and approved by local ethics committee and the UK Home Office.

### Animals

Transgenic Kir4.1-floxed mice (strain: B6.129-*Kcnj10^tm1Kdmc^*/J; Jackson Laboratory; strain number: 026826) of both sexes (male and female) and age between 2-4 months were used in this study. Heterozygous mice were imported and bred according to the provider’s instructions to establish a colony. Ear samples of each animal were genotyped at Transnetyx, and only the homozygous mice were used in the study. Mice were housed in individually ventilated cages in a 12-hour light and 12-hour dark lighting regime. Mice received food and water *ad libitum*.

#### Focal Kir4.1 conditional knockout mice

To generate focal Kir4.1 conditional knockout mice (Kir4.1-cKO), AAV-PHP.B2A-gfaABC1D-mCherry-IRES-Cre, an adeno-associated viral vector (AAV) expressing Cre recombinase under astrocyte specific promoter (gfaABC1D) was generated in-house and injected unilaterally into the hippocampus of adult Kir4.1-floxed mice. For further details, see ‘Surgeries’ section.

### AAV-Cre vector cloning, production and quantification

The pAAV-gfaABC1D-mCherry-IRES-Cre vector genome was generated by replacing the EF1α promoter of pAAV-EF1α-mCherry-IRES-Cre (Addgene #55632) with the astrocyte-specific gfaABC1D promoter. Vector genomes were packaged into PHP.B2A capsids (kindly provided by Prof. Viviana Gradinaru, California Institute of Technology) by triple-plasmid co-transfection of HEK293T cells (ATCC, #CRL-11268) with the vector genome plasmid, PHP.B2A capsid plasmid and pXX6.80 adenoviral helper plasmid (North Carolina Vector Core). AAV vectors were harvested from cells and culture medium, purified by iodixanol density-gradient ultracentrifugation, and concentrated in PBS containing 0.001% Pluronic F-68.^24^ Vector genome titres were determined by quantitative PCR using linearised plasmid standards. Viral purity was assessed by SDS-PAGE followed by SYPRO Ruby protein staining (Invitrogen).

### Surgeries

All surgical procedures were performed under aseptic conditions.

#### Animal preparations

For all surgeries, animals were weighed and anaesthetised with isoflurane in an induction chamber and positioned in a stereotaxic frame (David Kopf Instruments Ltd, USA) while maintained under isoflurane anaesthesia. Viscotears (Bausch+Lomb) was applied to the eyes to prevent drying. Analgesia was provided using buprenorphine (0.5 mg/kg, subcutaneous (s.c.); Buprevet, Virbac) and meloxicam (15 mg/kg; s.c.; Metacam, Boehringer Ingelheim). Saline (0.3 mL) was administered subcutaneously every 45 minutes depending on the surgery duration and again at the end of the procedure. Mice were monitored daily for at least five consecutive days following surgery.

#### AAV vector injections and subcutaneous transmitter implantation

The head was shaved and the skin disinfected with 50% (v/v) iodine. A midline incision was made to expose the skull and the connective tissue overlying the skull was gently removed.

##### AAV vector injection

A burr hole was drilled above the right hippocampus at the following coordinates relative to bregma (mm): AP, -2.3; ML, 2.0. To induce focal astrocyte-specific Kir4.1 knockout in the hippocampus, AAV-PHP.B2A-gfaABC1D-mCherry-IRES-Cre (injection volume, 1.8 µL; titre: 1.66x10^13^ vg/mL), was injected (DV: 1.25 and 1.75 mm from brain surface). Hereafter AAV-PHP.B2A-gfaABC1D-mCherry-IRES-Cre will be referred to as Ast-Cre viral vector. Control animals received pAAV-GFAP104-mcherry virus (injection volume, 1.8 µL; titre: 1x10^13^ vg/mL; Addgene, #58909-AAV5) hereafter referred to as control viral vector. Viral vector injections were performed using a 5 µL Hamilton syringe (Model 95, Hamilton Company, Switzerland) fitted with a 33-gauge needle and controlled by a microinjection pump (WPI Ltd., USA) at a rate of 150 nL/min. The virus was delivered at two depths: half the volume at 1.75 mm below the brain surface and, after a 5-minute pause, the remaining volume at 1.25 mm. After injection, the needle was slowly withdrawn with a 20-second pause every 0.2 mm to minimise backflow.

##### Electrode implantation

Following viral vector injection, a second incision was made along the shaved back of the animal. The skin and the underlying connective tissue were separated to create a pocket for a subcutaneous wireless transmitter (A3048, Open Source Instruments Inc., USA). The transmitter was placed in this pocket and the electrode wires were tunnelled subcutaneously to the skull. The recording wire was connected to a depth electrode (W-electrode, SDE-W, Open Source Instruments Inc.) and inserted through the burr hole at following coordinates relative to bregma (mm): AP, -2.3; ML, 2.0; DV, 1.25. The electrode was secured with a small drop of dental cement (Simplex Rapid, Associated Dental Products, UK). A second burr hole was drilled above the contralateral somatosensory cortex (AP, -1.0; ML, -3.0) to insert the reference wire, which was secured with a screw. An additional support screw was placed on the contralateral hemisphere. The exposed skull and the electrodes were secured with dental cement and the incision on the back was sutured.

#### AAV vector injections, headplate attachment, and craniotomy for acute awake head-fixed experiments

For acute awake head-fixed experiments, mice underwent two separate surgical procedures. In the first surgery, the head was shaved and the skin was disinfected with 50% (v/v) iodine, followed by a small midline incision to expose the skull. The connective tissue overlying the skull was gently removed. A burr hole was drilled to inject a viral vector to express channelrhodopsin in pyramidal neurons (pAAV-CaMKIIa-hChR2(H134R)-EYFP; injection volume, 0.5 µL; titre: 1x10^13^ vg/mL; Addgene, #26969-AAV9) into the hippocampus at the following coordinates relative to bregma (mm): AP, -2.3; ML, 2.0; DV, 1.3. This viral vector is hereafter referred to as AAV-CaMK2a-ChR2 viral vector. Following the injection, the incision was sutured and animals were allowed to recover for 14 days before the second surgery. In the second surgery, the skin over the skull was excised and the connective tissue was removed to expose the skull. The skull was cleaned and dried, and the Ast-Cre viral vector (injection volume, 0.5 µL; titre: 1.66x10^13^ vg/mL), was injected into the hippocampus at the same coordinates as the AAV-CaMK2a-ChR2 viral vector. All viral vector injections were performed at the rate of 150 nL/minute using 5 µL Hamilton syringe (Model 95 Hamilton Company, Switzerland) fitted with a 33-gauge needle and controlled by a microinjection pump (WPI Ltd., USA). A headplate (Model 9, Neurotar) was attached to the skull using Vetbond (1469SB, 3M) and further secured with dental cement. After the cement had fully cured, a craniotomy was performed to expose two areas: (a) a small (∼2×2 mm) craniotomy over the ipsilateral hippocampus (AAV injection side) for optogenetic stimulations and electrophysiological recordings, and (b) a burr hole contralateral to the main craniotomy for placement of a reference wire (silver/silver chloride). The exposed dura was covered sequentially with cortex buffer solution (mM: 125, NaCl; 4, KCl; 10, glucose; 10, HEPES; 2, CaCl2; 2, MgCl2. pH, 7.35), sterilised Sylgard 184 (∼200 μm thickness), a Kwik-Cast layer and a final layer of Kwick-Sil (WPI, UK). For the second surgery, dexamethasone (1 mg/kg, s.c.; Dexafast, Livisto) was administered in addition to the analgesics described in the ‘Animal preparations’ section. Mice were allowed to recover and monitored daily after surgery.

#### AAV vector injections and DC-coupled head-mount transmitter (DC-HMT) implantation

The head was shaved and the skin was disinfected with 50% (v/v) iodine. The skin over the skull was excised to expose the skull surface. The connective tissue overlying the skull was gently removed, and the skull was cleaned and dried.

##### AAV vector injection

Ast-Cre viral vector was injected unilaterally into the right hippocampus as described in the ‘AAV vector injection’ section under ‘AAV vector injection and subcutaneous transmitter implantation’ subheading.

##### Electrodes implantation and DC-HMT attachment

Following viral injection, four recoding electrodes from an electrode interface fixture (EIF; EIF8-XAAA; Open Source Instruments Inc., USA) were implanted at the following coordinates relative to bregma (mm): a) ipsilateral hippocampus (iHipp; AP, -2.3; ML, 2.0; DV, 1.25), b) ipsilateral somatosensory cortex (iSCtx; AP, -0.5; ML, 2.7), c) ipsilateral motor cortex (iMCtx; AP, 2.4; ML, 0.9), and d) contralateral somatosensory cortex (cSCtx; AP, -1.0; ML, -3.0). Of the four recording electrodes, one was a depth electrode inserted into the ipsilateral hippocampus, while the remaining three were wire electrodes secured with screws for recording from cortical regions. An additional burr hole was drilled to insert a reference wire into the cerebellum which was also secured with a screw. The exposed skull and all electrodes were sealed with dental cement. After the dental cement had cured, a wireless, battery-powered DC-HMT (A3040D3Z; Open Source Instruments Inc., USA) was connected to the EIF via the integrated Omnetics connectors. Animals were allowed to recover before being returned to their home cages.

### *Ex vivo* experiments

#### AAV vector injection

Ast-Cre viral vector was injected unilaterally into the right hippocampus as described in the ‘Viral injection’ section under the ‘AAV vector injection and subcutaneous transmitter implantation’ subheading. Following viral injection, mice were allowed to recover and maintained for 10-14 days before further experiments.

#### Hippocampal slice preparation

Transverse hippocampal slices (350 μm thick) from both hemispheres (AAV injected and non-injected sides) were prepared from Kir4.1-cKO mice 10-14 days after viral vector injection. The hippocampal slices from the non-injected hemisphere were used as controls. Hippocampal tissue was sliced in an ice-cold slicing solution containing (in mM): sucrose 75, NaCl 87, KCl 2.5, CaCl2 0.5, NaH2PO4 1.25, MgCl2 7, NaHCO3 25, and D-glucose 25. Slices were allowed to recover for 20 minutes at 34°C in the same solution and were then transferred to artificial cerebrospinal fluid containing (in mM): NaCl 119, KCl 2.5, NaH2PO4 1.25, MgSO4 1.3, CaCl2 2.5, NaHCO3 25, and D-glucose 11. All solutions were continuously bubbled with 95% O2 and 5% CO2, and osmolarity was adjusted to 298±3 mOsm.

#### Electrophysiological recordings

To assess synaptic transmission at CA3-CA1 synapses, extracellular recordings were obtained from the CA1 stratum radiatum in both control (non-injected hemisphere) and Kir4.1-cKO slices, with recordings in Kir4.1-cKO slices targeted to regions containing mCherry-labelled astrocytes. Hippocampal slices were transferred to a recording chamber mounted on an Olympus BX51WI upright microscope (Olympus, Tokyo, Japan) and superfused at 32-34°C and field excitatory postsynaptic potentials (fEPSPs) were recorded using a standard 2 MΩ glass pipette filled with perfusion solution. Synaptic responses were evoked by stimulating the Schaffer collateral fibres with a bipolar electrode placed in the stratum radiatum, at a distance of more than 200 μm from the recording site. Stimulation trains The stimulation intensity was set at approximately 50% of the maximal response and delivered at frequencies of 5, 20, and 50 Hz each for 5 seconds (pulse width: 200 µs).

#### Multiplexed 2PE imaging *ex vivo*

To assess extracellular potassium dynamics, hippocampal slices were incubated with the extracellular potassium ion biosensor GINKO2 (a gift from Prof. Robert Campbell; Addgene #177116)^25^ for 1 hour prior to recordings as previously described by Tyurikova *et. al*.^6^ 2PE acquisitions were performed using a Femto2D imaging system (Femtonics, Budapest) with laser at λ_x_^2P^=940 nm optimised for GINKO2 excitation, for selecting areas in the stratum radiatum with a bright GINKO2 signal (green channel) that also contained mCherry-positive astrocytes (red channel). For mCherry visualization, a second laser at λ_x_^2P^=1040 nm was used as detailed previously.^6^ For controls, slices with bright GINKO2 signal in the stratum radiatum of the non-injected hippocampus were selected. Once a suitable region was identified, frame scans of the selected area, containing 2-3 astrocytes, were acquired over a 20-second period. Fluorescence signals from the regions of interest (ROI) were analysed. Evoked changes in extracellular GINKO2 fluorescence were quantified as the ratio of peak signal intensity to baseline (dF/F0), using previously established routines for monitoring extracellular fluorescence transients.^26^

### Awake, head-fixed electrophysiology recordings and optogenetic stimulation

Mice implanted with headplates were habituated to the head-fixed experimental setup after at least 4 days of post-surgery recovery. Habituation was performed by placing the animals in the Neurotar frame for progressively increasing durations (15-60 min) over 3-4 days. Acute awake head-fixed experiments were performed 10-14 days after Ast-Cre viral vector injection. On the day of the experiment, mice were head-fixed in the Neurotar frame while allowing to move freely on the floating carbon fibre plate. Craniotomies were exposed by removing the Kwik-Sil, Kwik-Cast and Sylgard layers. The exposed dura was covered with cortex buffer solution.

Electrophysiological signals were recorded using graphene-based depth neural probes (gDNP).^12^ gDNPs are flexible probes capable of recording wide-band DC-coupled electrophysiological signals,, thereby enabling recording of both spreading depolarizations and seizure activity. Briefly, temporary stiffening of the flexible gDNP was achieved by back-coating with silk fibroin in a microstructured polydimethylsiloxane mould, and curing at 80 °C for 90 minutes followed by 10 minutes at room temperature (for further details, see Calia *et. al*).^12^ After curing, probes were removed from the mould and connected to printed circuit boards (PCB) connected to a custom g.RAPHENE amplifier (G.TEC Medical Engineering, GmbH, Austria) and interfaced with a computer. The gDNP connected to the PCB was lowered using a micromanipulator to just above the dura (∼0.1 mm lateral to the Ast-Cre viral vector injection site). The dura was punctured using a 26-gauge needle and the gDNP was inserted ∼1.6 mm into the brain. A silver/silver chloride refence wire was inserted into the contralateral motor cortex. A transfer curve was generated to determine the optimal gate voltage (Vgs) prior to initiating electrophysiological recording (sampling frequency: 9600 Hz).^12,27^

For optogenetic stimulation, an optic cannula (200 µm core diameter, 0.39 NA; CFMC12L20, Thorlabs) was positioned at the AAV-CaMK2a-ChR2 vector injection site, lowered to just above the dura, and inserted ∼1-1.2 mm into the brain. The cannula was coupled to a patch cable (200 µm core, 0.39 NA; M77L01, Thorlabs) via a mating sleeve (ADAF1, Thorlabs). The patch cable was connected to a fibre-coupled LED (470 nm; M470F3, Thorlabs) controlled by an LED driver (DC2200, Thorlabs) interfaced with a computer. Prior to the insertion, the optical power at the fibre tip was measured using a power meter (PM16-130; Thorlabs). For the stimulation intensity used in the experiments, the power measured at the fibre tip was 2.73 mW, corresponding to an estimated irradiance of 86.9 mW/mm². Optogenetic stimulation was delivered using Spike2 software (Cambridge Electronics Design limited, UK), which generated external trigger signals to control the LED via the DC2200 driver. Stimulation consisted of 10-second trains delivered at 5, 20, and 50 Hz with a pulse width of 10 ms. Electrophysiological responses were simultaneously recorded using gDNP. A minimum inter-stimulation interval of 15 minutes was maintained between stimulation trials. Each experiment was limited to a maximum duration of 90-100 minutes in accordance with project license and UK Home Office regulations. At the end of the experiment, mice were euthanised by intraperitoneal administration of sodium pentobarbital.

### Chronic *in vivo* telemetry recordings

For both subcutaneous transmitter (SCT) and DC-coupled head-mount transmitter (DC-HMT) recording experiments, mice were individually housed, and cages were placed within a Faraday enclosure. Intracranial EEG (iEEG) signals from the transmitters were received by antennas positioned adjacent to the cage and connected to an Octal data receiver and LWDAQ driver (Open Source Instruments Inc., USA), which interfaced with a computer. Continuous iEEG signals were acquired using LWDAQ software (Open Source Instruments Inc.) for 21 days following viral vector injection (day-0). Signals recorded using SCT and DC-HMT systems were sampled at 256 Hz and 512 Hz, respectively. In video-DC-HMT experiments, continuous video recordings were acquired using IP cameras (Microseven, USA) at 30 frames/second for the duration of the experiment. Video times were synchronised with the iEEG recording computer clock via the Windows time server to enable alignment with electrophysiological recordings. The DC-HMT was powered by a button battery (CR1225; RS Components) which was replaced once every 6 days.

### Western Blot

Mice were injected with Cre or control viral vector unilaterally into the right hippocampus as detailed in ‘AAV vector injection’ section under ‘AAV vector injection and subcutaneous transmitter implantation’ sub-heading. 21 days post-viral injection, mice were deeply anaesthetised (pentobarbital overdose i.p., Pentoject, Ecuphar, Belgium), decapitated, and the brains were immediately removed. The hippocampi were dissected and snap-frozen in the liquid nitrogen. Tissue lysate was prepared via homogenization straight from frozen using Micropestle (Eppendorf) in T-PER Tissue Protein Extraction Reagent (Thermo) supplemented with 1x Halt Protease and Phosphatase Inhibitor Cocktail (Thermo) according to the manufacturer’s instructions. Protein concentration was measured using the Pierce BCA Protein Assay Kit (Thermo) according to the manufacturer’s instructions. Western blotting was performed as previously described.^28^ Briefly, equal amount of each protein sample was resolved on a 4-15% Mini-PROTEAN TGX Precast Protein Gel (Bio-Rad) side-by-side with the Novex Sharp Pre-stained protein standard (Invitrogen). Subsequently, separated proteins on the gel were transferred onto the Immobilon-P polyvinylidene fluoride (PVDF) membrane (Millipore). The resulting blots were blocked in 5% milk in 0.1% PBS-Tween (v/v) for 1 hour at room temperature and then probed with Rabbit anti-Kir4.1 primary antibody (Proteintech, 12503-1-AP) diluted 1:1000 in 5% milk in 0.1% PBS-Tween for overnight at 4°C. The following day, blots were washed and incubated with Donkey anti-Rabbit IgG HRP (Abcam, ab16284) secondary antibody diluted 1:2000 in 5% milk in 0.1% PBS-Tween for 2 hrs at room temperature. HRPs on the immunoblots were enhanced using the SuperSignal West Pico Chemiluminescent Substrate (Thermo) according to the manufacturer’s instructions. The chemiluminescent images of membranes were captured using GeneGnome XRQ imaging system and GeneSys image acquisition software (Syngene). Next, the blots were stripped using Restore PLUS Western Blot Stripping Buffer (Thermo) according to the manufacturer’s instructions and then re-probed with Rabbit anti-GAPDH primary antibody (Cell Signalling Technology, 2118, 1:2500) as loading control. Densitometry analysis on images was performed using Fiji to quantify the intensity of each protein band, which was then normalized to the intensity of each corresponding GAPDH band, and the relative protein content was expressed as arbitrary units.

### Immunohistochemistry, microscopy and transduction volume quantification

Mice were deeply anaesthetised (pentobarbital (i.p.), Pentoject, Ecuphar, Belgium) and transcardially perfused with chilled 20 U/ml heparin (Sigma) in PBS followed by 4% PFA in PBS. Subsequently, the brains were immediately removed and post-fixed in 4% PFA in PBS at 4°C for 2-4 hrs. Fixed brains were transferred to 30% sucrose (Sigma) in PBS for cryo-protection until they sank and then frozen in O.C.T. compound (VWR) for cryo-sectioning. The brains were cryo-sectioned into 35 µm coronal sections of 12 series using a Leica CM1850 Cryostat.

Immunohistochemistry staining of free-floating brain sections was conducted as previously described.^28^ In brief, sections were blocked in 10% donkey serum (Abcam)/0.25% Triton X-100 (Sigma)/PBS at room temperature for 1 hour, then incubated with goat anti-mCherry (LSBio, LS-C204207) and rabbit anti-GFAP (Abcam, ab7260) primary antibodies diluted 1:1000 in 10% donkey serum/0.25% Triton X-100/ PBS for 40 hrs at 4°C. Subsequently, sections were washed 4 times with PBS and re-blocked in 10% donkey serum/0.25% Triton X-100/PBS at room temperature for 30 minutes. Sections were then incubated with donkey anti-Goat Alexa Fluor 594 (Invitrogen, A11058) and donkey anti-rabbit Alexa Fluor 488 (Invitrogen, A21206) secondary antibodies diluted 1:500 in 10% donkey serum/0.25% Triton X-100/PBS for 1 hour at room temperature. The sections were then mounted onto the Superfrost Plus microscope slides (VWR) with ProLong Gold Antifade Mountant with DAPI (Invitrogen) and sealed with nail polish.

Multi-channel fluorescence tiling images were captured using a Zeiss Observer invert widefield microscope, with 10x objective at 1024 x 1024 pixels resolution. Confocal images were acquired using a Zeiss LSM710 invert confocal microscope with the oil-immersion 63x objective, 2x scanner zoom, 956 x 956 pixels resolution and z-series of images taken at 0.13 µm intervals followed by deconvolution using Huygens (Scientific Volume Imaging) and 3D image processing by Imaris (Oxford Instruments). To quantify transduction volume, tiling images of phase contrast and red fluorescence channel were acquired using a Nikon Ti2 widefield fluorescence and brightfield microscope with 10x objective. All images were acquired using the same microscope parameters for each type of microscopy. Transduction volume of Cre recombinase labelled with red fluorescent mCherry signals in the hippocampi and cortices of both sides were quantified with FIJI as previously described.^27^ Briefly, the right and left hippocampi and cortices of each animal were manually selected as the regions of interest (ROIs) according to the Allen Mouse Brain Atlas (Atlas Thumbnails :: Allen Brain Atlas: Mouse Brain) and an intensity threshold consistent across the animals was applied. Pixels above the threshold were counted with total area in mm^2^ calculated based on pixel dimensions and magnification. The transduced area within each 35 µm slice was taken as a representation of the surrounding anterior-posterior 420 μm of tissue. The total transduction volume (mm^3^) was calculated as the summation of each 420 μm volume.

### Data analysis

#### Seizure detection

Seizure detection pipeline was adapted from the approach described by Wykes *et. al.* ^29^ Briefly, focal iEEG signals recorded from each animal using one-channel subcutaneous transmitters were analysed using the Neuroplayer Event Classifier tool in the LWDAQ software (Open Source Instruments). Recordings were segmented into 1-second and 4-second epochs, and analysed using six metrics: power, coastline, intermittency, coherence, asymmetry and spikiness. For seizure detection, representative 4-second seizure segments were added to Kir4.1-cKO seizure libraries and used to automatically screen all the 4-second iEEG segments. Segments with signal power exceeding a threshold, and exhibiting similarity to library seizure events were selected as candidate seizure events. The same threshold and similarity criteria were applied to all recordings. Candidate events were subsequently reviewed manually to confirm individual seizure occurrence and determine seizure duration (Supplementary Fig. 1A-C). For spike detection, representative 1-second single-spike containing iEEG segments were added to a Kir4.1-cKO spike library. All 1-second iEEG segments were screened using the Event Classifier based on coastline, intermittency, coherence, asymmetry and spikiness metrics (Supplementary Fig. 1D, E). For more details on the event classifier, consult the Event Classifier manual on the Open Source Instruments website https://www.opensourceinstruments.com/Electronics/A3018/Neuroplayer.html#Event%20Classifier.

#### DC shift

In acute awake head-fixed experiments, DC-shift during optogenetic stimulation was defined as the maximum voltage difference between the baseline period (10 seconds immediately preceding stimulation) and the stimulation period.

#### Power analysis

Spectral power was calculated using custom-written scripts in MATLAB with the built-in *pspectrum* function (frequency range: 0.001-100 Hz; frequency resolution: 2 Hz). Seizure power was normalised to the baseline power. Baseline was defined as a 10-second period without epileptiform activity (seizures, spikes, or spreading depolarization). For awake head-fixed experiments, baseline period was selected at least 50 seconds before the start of the onset of optogenetic stimulation.

#### Coastline analysis

Coastline measures were calculated for seizures and baseline periods using the Neuroplayer tool in LWDAQ software (Open Source Instruments). Baseline was defined as a 10-second period without epileptiform activity (seizures, spikes, or spreading depolarization). Seizure coastline values were normalised to baseline.

#### Seizure inter-event intervals

In the DC-HMT dataset, Seizures were identified manually. The inter-event intervals (IEI) were defined as the time between the end of one seizure and the onset of the subsequent seizure. For two consecutive inter-event-interval analysis (Fig. 6B), the first and second IEI value is normalised to the total sum of the two consecutive IEIs i.e., IEI_1_norm_ = IEI_Sz1-Sz2_ / (IEI_Sz1-Sz2_+ IEI_Sz2-SzSD_) and IEI_2_norm_ = IEI_Sz2-SzSD_ / (IEI_Sz1-Sz2_+ IEI_Sz2-SzSD_).

#### Postictal activity recovery analysis

Recovery of electrical activity following seizures was quantified by measuring changes in 1-40 Hz power. Power in 1-40 Hz band was calculated using custom-written MATLAB scripts with the built-in *psprectrum* function (frequency resolution: 2 Hz). The resulting power-time series was smoothed using moving-mean method with a 5-second window. Power values were normalised to the pre-seizure baseline and expressed as percentage change relative to baseline. Mean 1-40 Hz power was measured at eight pre-defined time points: pre-seizure baseline (bfr), during the seizure (Sz), and six postictal recovery periods (r1-r6). For acute awake head-fixed recordings, the first recovery period (r1) was defined as 100 seconds after electrographic seizure termination, with subsequent recovery periods separated by 100-second intervals. For chronic DC-coupled head-mount transmitter experiments, the first recovery period (r1) was defined as 50 seconds following electrographic seizure termination, with subsequent recovery periods separated by 50-second intervals. In all experiments, mean power was calculated within a 10-second window for the baseline and each recovery time point. Recovery time is defined as the time required for the 1-40 Hz power to reach 80% of the baseline and remain above this threshold continuously for at least 5 seconds. If this criterion was not met, the final analysed time point (r6) was assigned as the recovery time.

#### Postictal depression

Postictal depression was quantified as the percentage change in 1-40 Hz power during the 120-second period following electrographic seizure termination relative to the pre-seizure period. For principal component and logistic regression analyses, postictal depression was quantified as the ratio of 1-40 Hz power measured before seizure onset to that measured for 120 seconds after seizure termination.

#### Principal component analysis

Principal component analysis (PCA) was performed to determine whether seizure-alone and SD-associated seizures occupied distinct regions in the feature space. Three seizure-related features were included in the analysis: seizure power, seizure duration, and postictal depression. Variables were standardised (z-scored) to account for differences in scale. PCA was performed using GraphPad Prism (v10.6.1 GraphPad Software). Principal components were ranked according to their eigenvalues, and components explaining ≥75% of the total variance were retained for further analysis.

#### Multivariable Logistic Regression

A multivariable logistic regression model (event type ∼ seizure power + seizure duration + postictal depression) was used to evaluate whether electrophysiological features available in conventional high-pass-filtered recordings could retrospectively identify seizures associated with SD. Predictor variables included seizure power (25-80 Hz), seizure duration, and postictal depression. Model discrimination was evaluated using the area under the receiver operating characteristic (ROC) curve (AUC), and goodness of fit was assessed using the log-likelihood ratio test. Odds ratios (OR) with 95% confidence intervals were calculated to quantify the association between each predictor variable and classification as an SD-associated seizure event. Analyses were performed in GraphPad Prism (v10.6.1), with statistical significance set at *P* < 0.05.

#### Behavioural analysis

Behavioural analysis was performed for 12 events per group (seizures-alone and seizures with spreading depolarization). Only events with clear video recordings showing unobstructed animal behaviour were included. Animal behaviour was assessed during both seizure and the immediately following postictal period.

Seizure behaviour was scored using a modified Racine scale comprising stages 1-6: (1) behavioural arrest, intense mastication and facial movements such as blinking, repetitive ear movements, (2) head nodding, backward movements, (3) unilateral forelimb clonus, body jerks, (4) bilateral forelimb clonus, (5) stage 4 seizure with loss of postural tone, and (6) jumping or repetitive falling, or running around the cage. Time spent in each Racine stage was quantified and expressed as a percentage of the total duration of each seizure.

For postictal behavioural analysis, the first 120 seconds immediately following seizure termination was analysed. Postictal behaviour was classified into three categories: (1) exploring, (2) behavioural arrest, and (3) abnormal behaviour, including loss of posture, intermittent clonus, falling, or movement with an altered gait. Time spent in each behavioural category was quantified and expressed as a percentage of the total analysed period for each event (120 seconds).

#### Ordinal Logistic Regression

Ordinal logistic regression analysis was performed to assess the relationship between seizure severity measures and postictal behavioural impairment. Postictal behavioural severity was treated as an ordinal dependent variable with three ordered levels: mild (exploratory behaviour), moderate (behavioural arrest), and severe (abnormal behaviour). For each seizure, postictal behavioural severity was assigned according to the behavioural category in which the animal spent the greatest proportion of postictal time. Maximum seizure Racine scores and seizure power (25-80 Hz) were included as predictor variables. Models were fitted using proportional odds ordinal logistic regression, and predictor significance was assessed using likelihood ratio tests. Odds ratios (ORs) with 95% confidence intervals (CIs) were calculated from model coefficients. Analyses were performed in Jamovi software (v2.6.44), with statistical significance set at *P* < 0.05.

#### Exclusion criteria

For inter-event interval analysis, events with intervals exceeding 720 mins (12 hours) were excluded. One seizure-associated spreading depolarization event followed by SUDEP was observed and excluded from postictal depression, principal component and regression analyses.

### Statistics

Statistical analyses were performed using GraphPad Prism (v10.6.1). Normality of datasets was assessed using the Shapiro-Wilk test. Data are presented as mean±SEM unless otherwise stated, and *n* values represent the number of data points with corresponding units. Comparisons between two groups were performed using paired or unpaired t-tests, as appropriate. When assumptions of equal variance were violated, Welch’s t-test was used for independent group comparisons. For non-normally distributed data, the Wilcoxon matched-pairs signed rank test or Mann-Whitney U test was used for paired or unpaired comparisons, respectively. For comparisons involving more than two groups, analysis of variance (ANOVA) with appropriate *post hoc* corrections was performed. Two-way ANOVA followed by Tukey’s multiple comparisons test was used for the analysis of DC shift (*ex vivo* and awake head-fixed datasets), GINKO2 amplitude (*ex vivo* dataset), and postictal behavioural measures (video-iEEG dataset). Two-way repeated-measures ANOVA followed by Tukey’s multiple comparisons test was used for analysis of viral vector transduction volume and activity recovery in the awake head-fixed dataset. One-way repeated-measures ANOVA followed by Tukey’s multiple comparisons test was used to assess differences in the power of successive seizures and seizures associated with spreading depolarization. For activity recovery analysis in the DC-HMT dataset, a linear mixed-effects model fitted using restricted maximum likelihood was applied to account for the unbalanced dataset, followed by Tukey’s multiple comparisons test. A *P* value < 0.05 was considered statistically significant.

### Data availability

The data that support the findings of this study are available from the corresponding author, upon request.

## Results

### Astrocytic Kir4.1 loss impairs hippocampal potassium buffering in the adult brain slices

To investigate the consequences of focal astrocytic Kir4.1 loss in an otherwise intact adult neural network, we conditionally deleted Kir4.1 within the hippocampus of adult Kir4.1-floxed mice by unilateral injection of a Cre recombinase-expressing viral vector under an astrocyte-specific promoter (Fig. 1A). This approach enabled investigation of acquired astrocytic potassium buffering dysfunction while avoiding developmental confounds. Immunofluorescence confirmed robust expression of Cre-mCherry in GFAP-positive astrocytes within the injected hippocampus (Fig. 1B). Western blot analysis further demonstrated a marked reduction in Kir4.1 protein levels in the injected hippocampus compared to controls (Fig. 1C), confirming effective focal reduction of astrocytic Kir4.1 expression.

**Figure 1.**
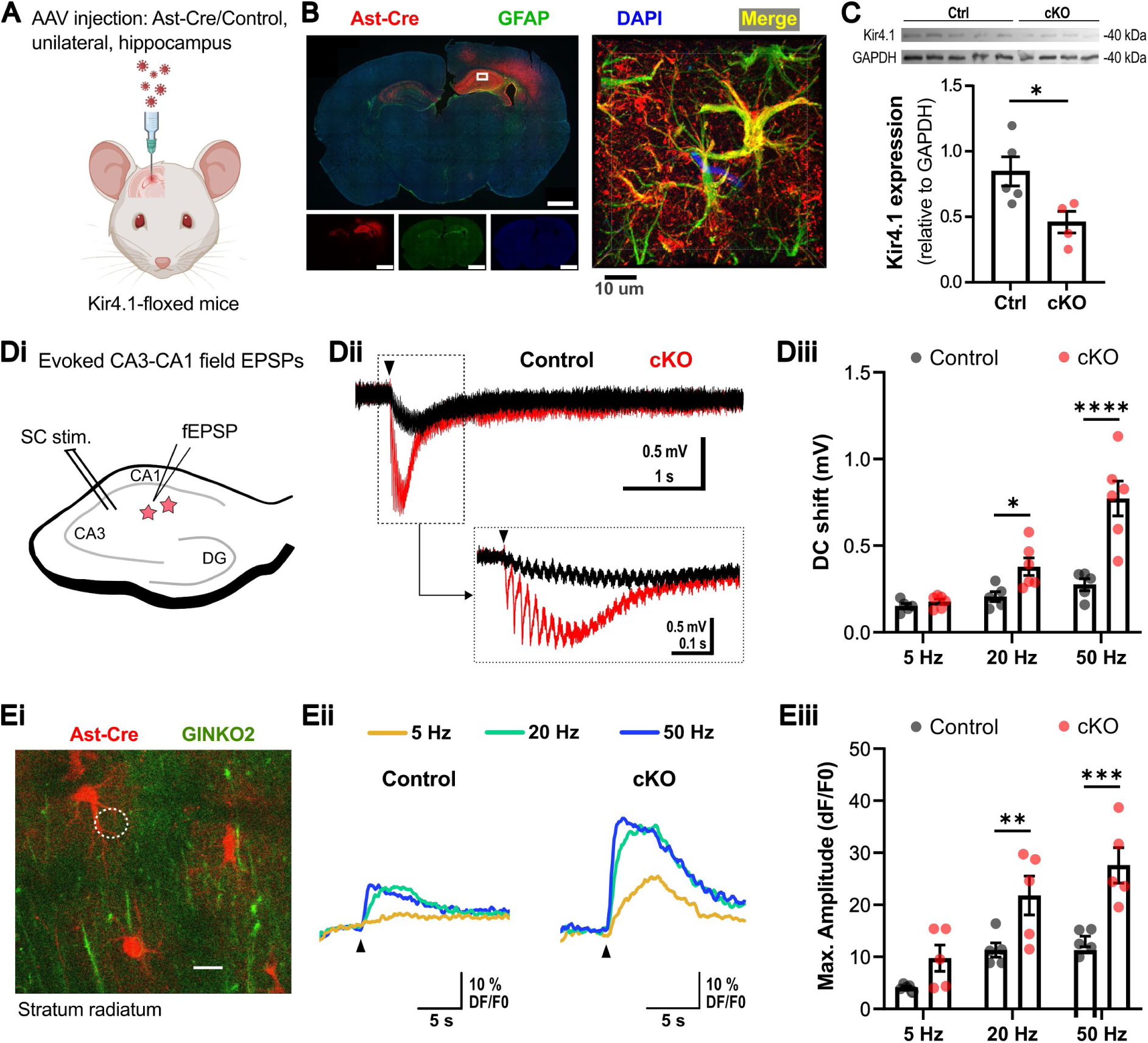
Astrocyte Kir4.1 loss impairs potassium buffering. (**A**) Experimental schematic showing unilateral, focal Ast-Cre or Control viral vector injection into the right hippocampus of adult Kir4.1-floxed mice. (**B**) Widefield fluorescence images show colocalisation (yellow) of Ast-Cre viral vector-tagged mCherry (red) and GFAP (green), with DAPI nuclear staining (blue). *Top-left*: representative whole-brain section from a Kir4.1-cKO mouse (scale bar, 1 mm). *Bottom-left*: corresponding individual channels (Ast-Cre, GFAP, and DAPI; scale bars, 2 mm). Right: higher-magnification view of the boxed region in the *top-left* image. (**C**) Detection of lower expression of Kir4.1 protein in Ast-Cre injected tissue (cKO) by western blot analysis. Controls: 0.85±0.11, *n* = 5 mice; cKO: 0.46±0.08, *n* = 4 mice; \**P* < 0.05. (**D**) *Di*, schematic of hippocampal slice showing Schaffer collateral stimulation and fEPSP recording in CA1 stratum radiatum. *Dii*, representative traces showing larger stimulation-evoked DC shifts in Kir4.1-cKO slices following Schaffer collateral stimulation (arrowhead, 50 Hz-5 s). *Diii.* DC shift magnitude increased with stimulation frequency and was significantly greater in Kir4.1-cKO slices. Two-way ANOVA, *F*(2, 27) = 23.39, *P* < 0.0001; Tukey’s multiple comparisons test, Control vs cKO: 5 Hz, *P* = 0.54; 20 Hz, \*\*\**P* < 0.0003; 50 Hz, \*\*\*\**P* < 0.0001. For mean±SEM and all comparisons, see Supplementary Table 1. (**E**) *Ei*, representative image of Ast-Cre–transfected astrocytes (red) and the extracellular potassium sensor GINKO2 (green) in a hippocampal slice. *Eii*, electrical stimulation (black arrow heads) of Schaffer collaterals induced larger GINKO2 fluorescence resposnses in cKO. *Eiii*, GINKO2 response amplitude (ΔF/F₀) increased with stimulation frequency and was significantly greater in Kir4.1-cKO slices. Two-way ANOVA, *F*(2, 24) = 16.31, *P* < 0.0001; Tukey’s multiple comparisons test, Control vs cKO: 5 Hz, *P* = 0.1137; 20 Hz, \*\**P* = 0.0053; 50 Hz, \*\*\**P* = 0.0002. For mean±SEM and all comparisons, see Supplementary Table 2.

Following confirmation of focal astrocytic Kir4.1 loss, we next examined the functional consequences for hippocampal potassium buffering and synaptic responses in acute brain slices. Schaffer collateral stimulation evoked a field response in the CA1 stratum radiatum accompanied by a negative DC shift consistent with activity-dependent extracellular potassium accumulation (Fig. 1Di, Dii). In slices from Kir4.1 knockout mice, these DC shifts were similar to such shifts in the controls at low frequency stimuli (5 Hz), thus confirming comparable stimulation conditions, yet become significantly larger at higher frequencies, indicating impaired potassium clearance from the extracellular space. The magnitude of the DC shift increased progressively with stimulation frequencies (20 and 50 Hz), consistent with enhanced activity-dependent potassium accumulation in Kir4.1-deficient tissue (Fig. 1Diii; Supplementary Table 1).

To directly measure stimulation-evoked changes in extracellular potassium, we employed the fluorescent extracellular potassium sensor GINKO2.^6,25^ Schaffer collateral stimulation evoked significantly larger fluorescent transients in Kir4.1 knockout slices compared to controls, with prolonged recovery kinetics following stimulation (Fig. 1Ei, Eii). Similar to the DC recordings, GINKO2 fluorescence signal amplitude increased as a function of stimulation frequency (Fig. 1Eiii; Supplementary Table 2). Together, these findings demonstrate that focal astrocytic Kir4.1 loss impairs activity-dependent extracellular potassium buffering within the adult hippocampus.

Given the central role of extracellular potassium regulation in controlling neuronal excitability, we next investigated whether focal loss of astrocytic Kir4.1 in the adult hippocampus is sufficient to drive spontaneous seizures *in vivo*.

### Focal astrocytic Kir4.1 loss induces spontaneous seizures

Ast-Cre viral vector was injected unilaterally into the hippocampus of adult Kir4.1-floxed mice to induce focal astrocytic Kir4.1 loss, and a subcutaneous wireless transmitter was implanted for chronic intracranial EEG (iEEG) recordings (Fig. 2A). Histological analysis confirmed robust viral vector transduction throughout the injected hippocampus with minimal spread to the ipsilateral cortex or contralateral hemisphere (Fig. 2B; Supplementary Table 3).

**Figure 2.**
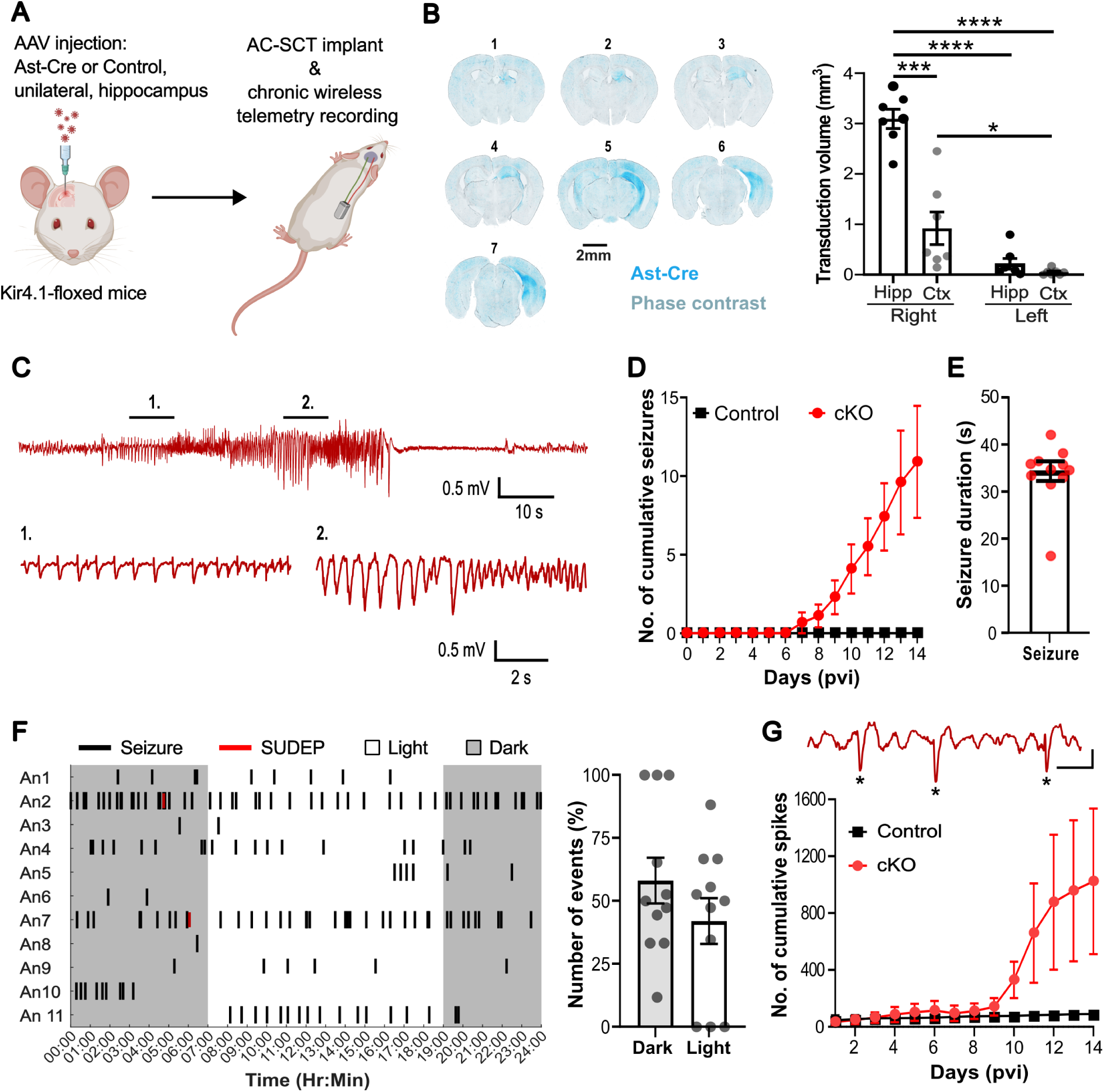
Focal Kir4.1 knockout in hippocampal astrocytes induce spontaneous seizures. (**A**) Experimental schematic showing unilateral hippocampal injection of Ast-Cre or control viral vectors and implantation of a subcutaneous AC-coupled wireless telemetry transmitter in Kir4.1-floxed mice. (**B**) *Left*: Representative rostro-caudal brain sections showing Ast-Cre expression (blue) in Kir4.1-cKO brain. *Right*: Quantification of transduction volume demonstrating a significant transfection of the injected (right) hippocampus. Two-way ANOVA, *F*(1, 6) = 43.14, *P* < 0.001; \**P* = 0.0256, \*\*\**P* = 0.002, \*\*\*\**P* < 0.0001. For mean±SEM and all comparisons, see Supplementary Table 3. (**C**) Representative hippocampal iEEG recording of a spontaneous seizure in a Kir4.1-cKO mouse, with expanded views of seizure onset (#1) and the later clonic phase (#2). (**D**) Cumulative seizure frequency plot show the development of spontaneous seizures around 7 days post-virus injection (pvi) in Kir4.1-cKO mice. No seizures were recorded in controls. 14 days pvi: cKO, 10.90±3.57 seizures, *n* = 11 mice; controls, *n* = 2 mice. (**E**) Seizure duration: 34.34±2.08 seconds, *n* = 11 mice. (**F**) *Left:* Circadian raster plot of seizure occurrence across light and dark phases. *Right:* Seizure distribution did not differ significantly between phases (dark, 58.02 ± 9.09%; light, 41.98 ± 9.09%; paired t-test, *P* = 0.3983; *n* = 11). (**G)** Cumulative spike frequency plot showing increased interictal spike occurrence beginning ∼7 days after Ast-Cre injection. 14 days pvi: cKO, 1023.31±511.60 spikes, *n* = 10 mice; controls, 90.50±4.50 spikes, *n* = 2 mice. Top: Representative hippocampal iEEG trace from Kir4.1-cKO showing spontaneous spikes (*). Scale bars: 0.5 s, 1 mV.

Spontaneous recurrent seizures were recorded from the ipsilateral hippocampus and identified using a semi-automated seizure detection pipeline (Fig. 2C; Supplementary Fig. 1). Seizures emerged approximately 7 days after Ast-Cre viral vector injection and progressively increased over time (Fig. 2D; Supplementary Fig. 2). Seizure activity remained detectable throughout the available recording period of up to 21 days, with seizure frequency peaking at 11.76±2.42 days post-injection (mean ± SD; Supplementary Fig. 2). At 14 days post-injection, Kir4.1-cKO mice exhibited a cumulative seizure frequency of 10.90±3.57 events (*n* = 11 mice), whereas no seizures were detected in control animals (*n* = 2 mice). Individual seizures had a mean duration of 34.34±2.08 s (Fig. 2E). These findings demonstrate that focal astrocytic Kir4.1 loss within the adult hippocampus is sufficient to drive spontaneous seizures.

To determine whether seizure occurrence followed a circadian pattern, seizure timing was analysed across light and dark phases. No significant difference in seizure distribution was observed between phases (Fig. 2F; Supplementary Fig. 2), indicating that seizures occurred throughout the circadian cycle. During chronic recordings, sudden unexpected death in epilepsy (SUDEP) was observed in 2/11 Kir4.1-cKO mice, with both events occurring during the dark phase (Fig. 2F).

In addition to spontaneous seizures, Kir4.1-cKO mice developed a marked increase in interictal spike activity beginning approximately 7 days after Ast-Cre viral vector injection (Fig. 2G). The progressive emergence of interictal epileptiform activity is consistent with increasing network hyperexcitability following focal astrocytic Kir4.1 loss. Together, these findings demonstrate that focal disruption of astrocytic Kir4.1 function within the adult hippocampus is sufficient to generate spontaneous epileptic activity *in vivo*.

### Astrocytic Kir4.1 loss increases susceptibility to evoked seizures and seizure-associated spreading depolarizations

Chronic wireless recordings demonstrated that focal astrocytic Kir4.1 loss is sufficient to generate spontaneous seizures *in vivo*. However, the telemetry system incorporated a high-pass filter (0.2 Hz) that prevents detection of spreading depolarizations (SDs). Given the emerging role of seizure-associated SDs in postictal dysfunction and SUDEP,^18,22^ we next sought to determine whether astrocytic Kir4.1 loss lowers the threshold for seizure induction and promotes seizure-associated spreading depolarizations.

To address this, we combined optogenetic activation of hippocampal pyramidal neurons with full-bandwidth DC-coupled electrophysiological recordings using graphene-based depth probes in awake, head-fixed mice (Fig. 3A). Graphene micro-transistor arrays provide stable full-bandwidth DC-coupled recordings, enabling concurrent high-fidelity detection of both seizures and spreading depolarizations.^12,27^ Experiments were performed 10-14 days after Cre or control viral vector injection, corresponding to the period of maximal seizure frequency observed during chronic recordings.

**Figure 3.**
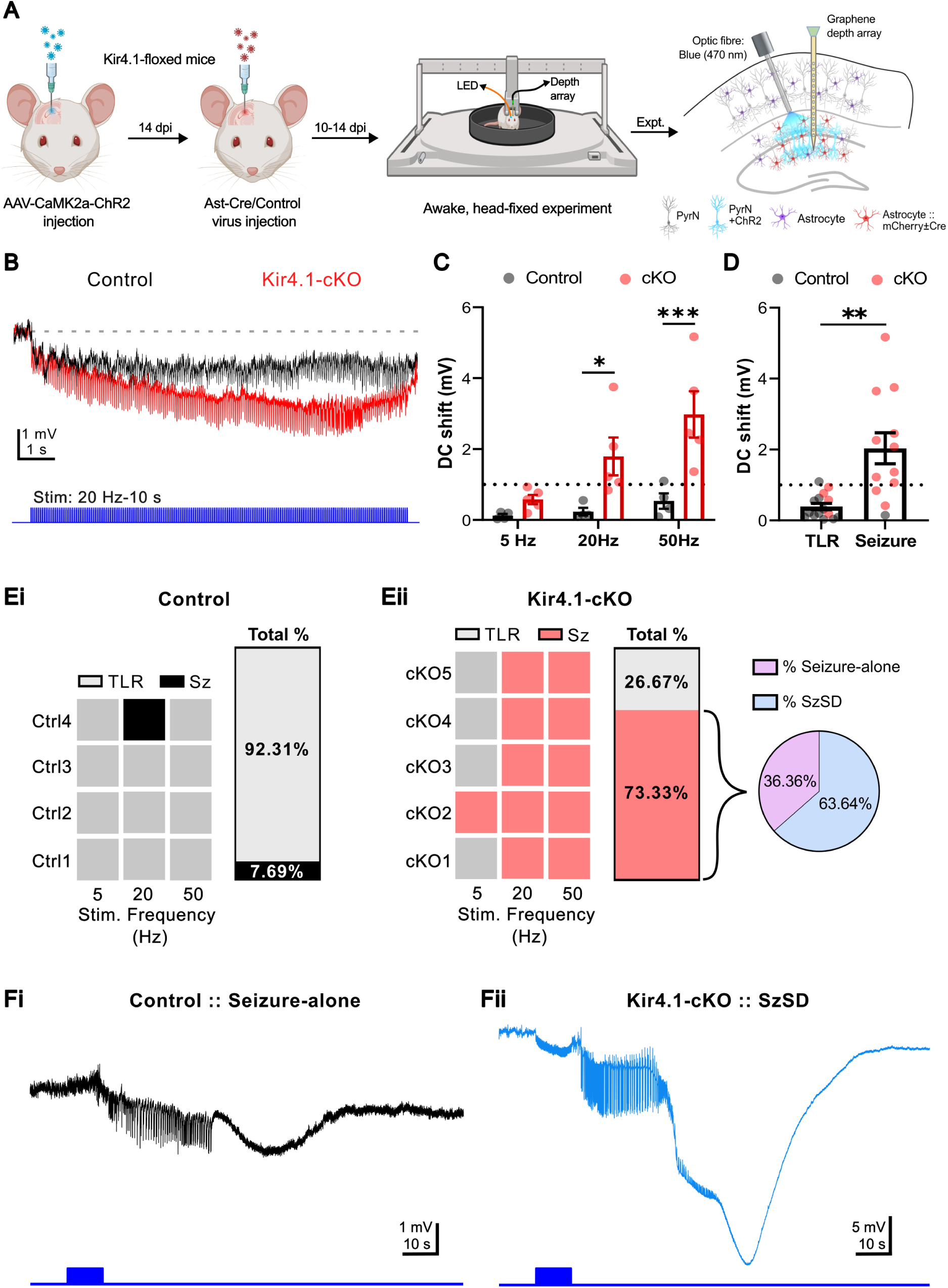
Optogenetic stimulation of pyramidal neurons triggers seizures and seizures temporally associated with spreading depolarizations in awake head-fixed Kir4.1-cKO mice. (**A**) Experimental schematic showing hippocampal injection of AAV-CaMK2α-ChR2, followed 14 days later by Ast-Cre or control viral vector injection and headplate implantation. Awake, head-fixed recordings were performed 10-14 days after Ast-Cre or control viral vectors injection using an optical fibre for optogenetic stimulation and a graphene-based depth probe for full-band LFP recordings. (**B**) Representative full-band LFP traces showing time-locked responses to optogenetic stimulation, with larger DC shifts in Kir4.1-cKO mice. (**C**) Optogenetically evoked DC shift magnitude increased with stimulation frequency and was significantly greater in Kir4.1-cKO mice. Two-way ANOVA: F (2, 21) = 5.99, *P* = 0.0087; Tukey’s multiple comparisons test, Control vs cKO: 5 Hz, *P* = 0.44; 20 Hz, \**P* = 0.01; 50 Hz, \*\*\**P* = 0.0004. For mean±SEM and all comparisons, see Supplementary Table 4. (**D**) Large stimulation-evoked DC shifts were associated with seizure generation. DC shift amplitude: Time-locked response (TLR), 0.39±0.09 mV, *n* = 15 trials; seizure, 2.04±0.44 mV, *n* = 12 trials; unpaired t-test, \*\**P* = 0.0032. (**E**) Optogenetic stimulation (5, 20, and 50 Hz) revealed increased seizure susceptibility in Kir4.1-cKO mice. *Ei*, control mice exhibited a single seizure during 20 Hz stimulation (1/13 trials; 7.69%), with the remaining trials producing TLRs (12/13; 92.31%). *Eii*, cKO mice exhibited seizures in 11/15 trials (73.33%) and TLRs in 4/15 trials (26.67%). Of the optogenetically induced seizures in cKO mice, 7/11 (63.64%) were associated with spreading depolarization (SzSD) and 4/11 (36.36%) occurred without SD. (**F**) Representative LFP responses to optogenetic stimiulation (20 Hz, 10 s). *Fi*, seizure-alone in control; *Fii,* SzSD in Kir4.1-cKO.

Optogenetic stimulation evoked reliable time-locked electrophysiological responses in both groups. However, Kir4.1-cKO mice exhibited markedly enhanced DC responses during stimulation compared with controls (Fig. 3B). The magnitude of these DC responses increased as a function of stimulation frequency and was significantly greater in Kir4.1-cKO mice (Fig. 3C; Supplementary Table 4). While responses to 5 Hz stimulation were comparable between groups, stimulation at 20 Hz and 50 Hz produced substantially larger DC shifts in Kir4.1-cKO mice, consistent with the impaired potassium buffering observed *ex vivo* (see Fig. 1Diii). Importantly, stimulation trials that progressed to seizures were associated with significantly larger DC shifts than trials producing only time-locked responses (unpaired t-test, *P* < 0.05; Fig. 3D). Across all stimulation trials, only a single seizure was observed in control animals (1/13 trials, 7.69%), whereas seizures occurred in the majority of trials in Kir4.1-cKO mice (11/15 trials, 73.33%; Fig. 3E). Notably, 7 of the 11 seizures recorded in Kir4.1-cKO mice were accompanied by spreading depolarizations, whereas no SD events were detected in control animals (Fig. 3E, F). The probability of both seizure generation and seizure-associated SDs increased with stimulation frequency in Kir4.1-cKO mice (Supplementary Fig. 3). Together, these findings demonstrate that focal astrocytic Kir4.1 loss lowers the threshold for seizure induction and strongly promotes seizure-associated spreading depolarizations.

To determine whether seizures associated with spreading depolarizations exhibit distinct electrophysiological and postictal characteristics, we compared spectral power and postictal recovery dynamics between seizures occurring with and without SDs (Fig. 4). Seizures associated with SDs exhibited significantly greater power within the beta (12-25 Hz), gamma (25-48 Hz), and high-gamma (52-98 Hz) frequency bands compared with seizures alone (Fig. 4A-C). In addition, seizures followed by SDs were associated with a markedly prolonged suppression of postictal electrographic activity (Fig. 4Di; Supplementary Table 5). Recovery of network activity was significantly delayed in seizures associated with SDs compared with seizures occurring in isolation (*P* < 0.05; Fig. 4Dii). Although there was a trend for seizures associated with SDs to be shorter in duration, this difference was not statistically significant between groups (Fig. 4E).

**Figure 4.**
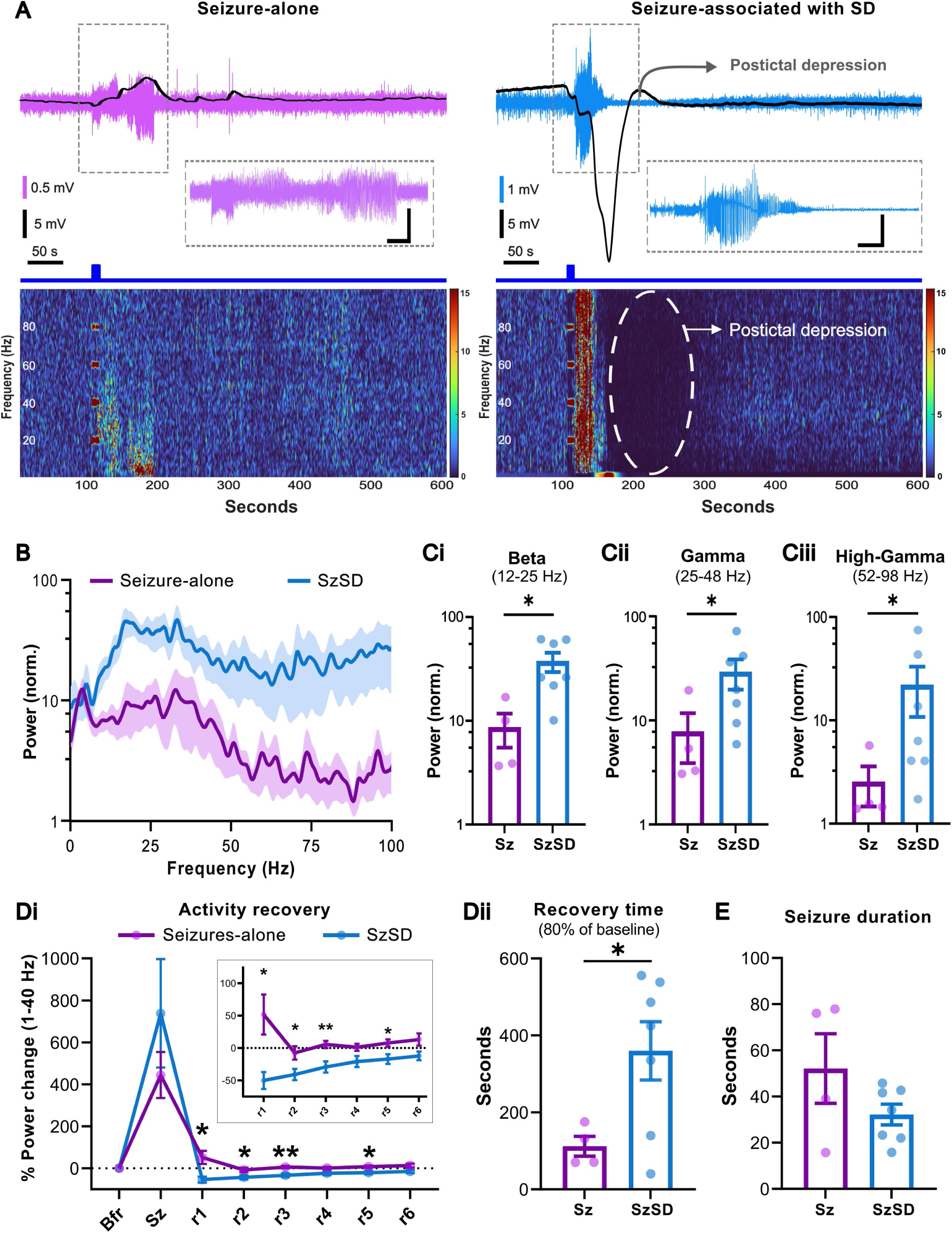
Seizures associated with spreading depolarizations show higher power and longer postictal recovery. (**A**) Representative seizure-alone (Sz; left) and seizure associated with SD (SzSD; right) events evoked by optogenetic stimulation (20 Hz, 10 s) in Kir4.1-cKO mice, with corresponding spectrograms. Band-pass filtered (1-100 Hz) traces are shown in magenta (Sz) and blue (SzSD), and DC (<0.1 Hz) traces in black. Spectrogram colour bars indicate normalised power. (**B**) Power spectral profiles of Seizure-alone and SzSD events. SzSDs exhibited greater power in the beta (12-25 Hz), gamma (25-48 Hz), and high-gamma (52-98 Hz) frequency bands. Solid lines and shaded regions represent mean±SEM. (**C**) Quantification of seizure power (normalised) showing significantly greater power during SzSD events (*n* = 7) compared with Seizure-alone (Sz) events (*n* = 4). *Ci*, beta band: Sz, 8.62±3.12; SzSD, 37.35±7.94; unpaired t-test, \**P* = 0.01; *Cii*, gamma band: Sz, 7.79±3.91; SzSD, 29.15±9.35; Mann Whitney test, \**P* = 0.04; *Ciii*, high-gamma: Sz, 2.53±1.06; SzSD, 21.90±11.13; Mann Whitney test, \**P* = 0.04. (**D**) Postictal recovery was prolonged following SzSD events. *Di*, percentage power change before, during, and after seizures (r1-r6), showing greater postictal depression and delayed recovery after SzSD. Two-way RM ANOVA, *F*(1.013, 9.115) = 10.95, *P* = 0.008; Tukey’s multiple comparisons test, Sz vs SzSD: r1, r2, and r5, \**P* < 0.05; r3, \*\**P* = 0.005. For mean±SEM and all comparisons, see Supplementary Table 5. *Dii*, Activity recovery time: Sz, 112.02±25.56 seconds, *n* = 4; SzSD, 360.10±75.84 seconds, *n* = 7; Welch’s t-test, \**P* = 0.0166. (**E**) Seizure duration: Sz, 52.14±15.09 seconds, *n* = 4; SzSD, 32.20±4.47 seconds, *n* = 7; Welch’s t-test, *P* = 0.2822.

Together, these findings demonstrate that seizure-associated SDs profoundly alter postictal network dynamics and are associated with prolonged suppression of electrographic recovery following seizures.

### Continuous DC-coupled telemetry reveals spontaneous seizures associated with spreading depolarizations in Kir4.1-cKO mice

Conventional chronic telemetry systems are unable to reliably capture DC shifts associated with spreading depolarizations (SDs), as recordings are typically high-pass filtered above 0.2 Hz. To overcome this limitation, we employed a novel DC-coupled head-mounted transmitter (DC-HMT) platform optimised for stable chronic full-bandwidth recordings in freely moving mice, enabling simultaneous detection of seizures and spreading depolarizations with minimal signal drift over prolonged recordings (Supplementary Fig. 4 and 5).

**Figure 5.**
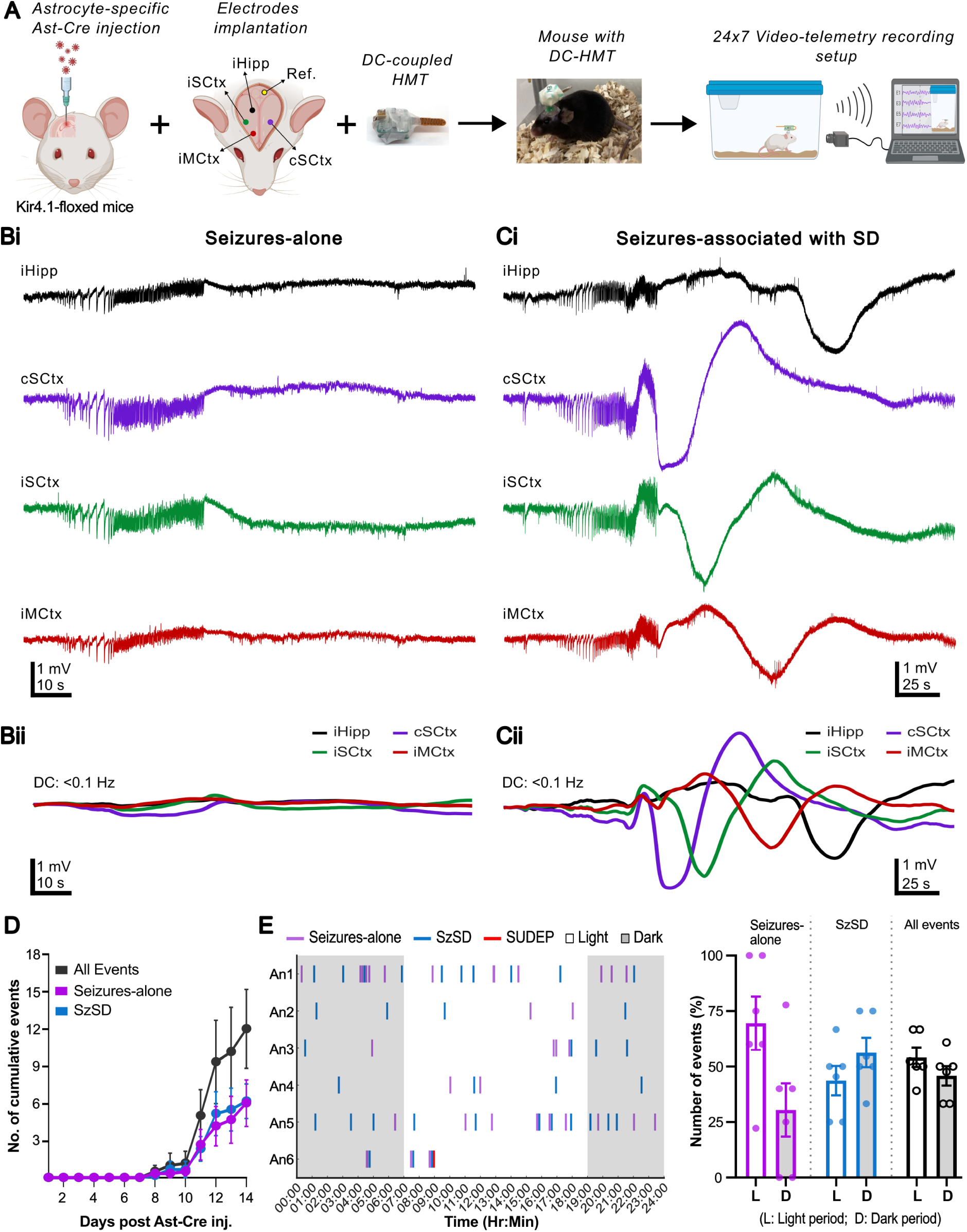
Chronic DC-coupled telemetry reveals spontaneous seizures and spreading depolarizations in freely moving Kir4.1-cKO mice. (**A**) Experimental schematic showing unilateral hippocampal Ast-Cre injection in Kir4.1-floxed mice, followed by electrode implantation, attachment of a DC-coupled head-mount transmitter (HMT), and continuous 24/7 video-telemetry recording in freely moving mice. (**B**) Representative seizure-alone events. *Bi*, full-bandwidth iEEG recordings. *Bii*, corresponding DC-band filtered traces (<0.1 Hz; overlaid). (**C**) Representative seizures associated with spreading depolarization (SzSD). *Ci*, full-bandwidth iEEG recordings. *Cii*, corresponding DC-band filtered traces (<0.1 Hz; overlaid). (**D**) Cumulative event plot showing the development of spontaneous seizure-alone and SzSD events ∼7–8 days after Ast-Cre injection. At 14 days post-virus injection (pvi), mice exhibited 5.83±1.89 seizure-alone events and 6.00±1.46 SzSD events (all events combined, 11.83±3.26). Event frequencies did not differ between seizure-alone and SzSD events (paired t-test, *P* = 0.8560; *n* = 6 mice). (**E**) *Left*, circadian raster plot showing the temporal distribution of seizure-alone (purple), SzSD (blue), and SzSD followed by death (SUDEP) (red) events over 14 days post-Ast-Cre injection. *Right*, seizure occurrence did not differ significantly between light and dark phases (*n* = 6 mice). Seizure-alone: light phase, 69.54±11.99%; dark phase, 30.46±11.98% (paired t-test, *P* = 0.1639); SzSD: light phase, 43.69±6.60%; dark phase, 56.31±6.61% (paired t-test, *P* = 0.3832); All events (seizure-alone and SzSD combined): light phase, 54.14±4.38%; dark phase, 45.86±4.37% (paired t-test, *P* = 0.3882).

To investigate whether spontaneous seizures in Kir4.1-cKO mice were associated with SDs, we performed continuous 24/7 video-iEEG recordings in six Kir4.1-cKO mice using DC-HMT devices (Fig. 5A). Spontaneous generalised seizures were detected in all six Kir4.1-cKO mice across all recording sites, including the ipsilateral hippocampus (iHipp), ipsilateral somatosensory cortex (iSCtx), ipsilateral motor cortex (iMCtx), and contralateral somatosensory cortex (cSCtx). These events comprised both seizures occurring in isolation (seizures-alone) and seizures associated with SDs (SzSD) (Fig. 5B,C). Surprisingly, in all the seizures associated with SDs, spreading depolarizations were first detected at a cortical recording site before subsequently appearing in other regions, consistent with propagation between cortical and hippocampal regions. However, due to the limited spatial sampling of recording electrodes, the precise site of SD initiation could not be determined. Both seizure-alone and seizure-associated SD events emerged approximately 7 days after Ast-Cre viral vector injection and continued throughout the recording period, occurring at broadly similar frequencies (Fig. 5D). We examined the temporal distribution of spontaneous seizure events across the light-dark cycle. Analysis of event timing revealed no clear circadian distribution for seizure-alone events, seizures associated with SDs, or all seizure events combined (Fig. 5E). During the recording period, a single SUDEP event was observed, occurring during the light phase.

### Spontaneous seizures associated with spreading depolarization exhibit distinct electrographic signatures and strong postictal depression

Following the identification of spontaneous seizures associated with SDs in Kir4.1-cKO mice, we examined whether these events emerged within temporally clustered seizure activity. Although seizures and seizure-associated SDs were otherwise distributed throughout the recording period without an obvious temporal pattern, we identified eight instances across five mice in which recurrent seizure-alone events occurred with progressively shortening inter-seizure intervals, culminating in a seizure associated with SD (Fig. 6A). Compared with preceding seizures within the same cluster, seizures associated with SDs exhibited greater electrographic power, longer duration, and increased coastline measures, consistent with progressive escalation of network activity prior to SD generation (Fig. 6B-D; Supplementary Fig. 6). Analysis of all recorded spontaneous events further revealed that seizures associated with SDs were followed by significantly longer intervals before the subsequent seizure compared with seizure-alone events (Fig. 6E). In addition, seizure-associated SDs produced markedly stronger and more prolonged postictal suppression of electrographic activity, with delayed recovery towards baseline network activity levels (Fig. 6F-H; Supplementary Table 6).

**Figure 6.**
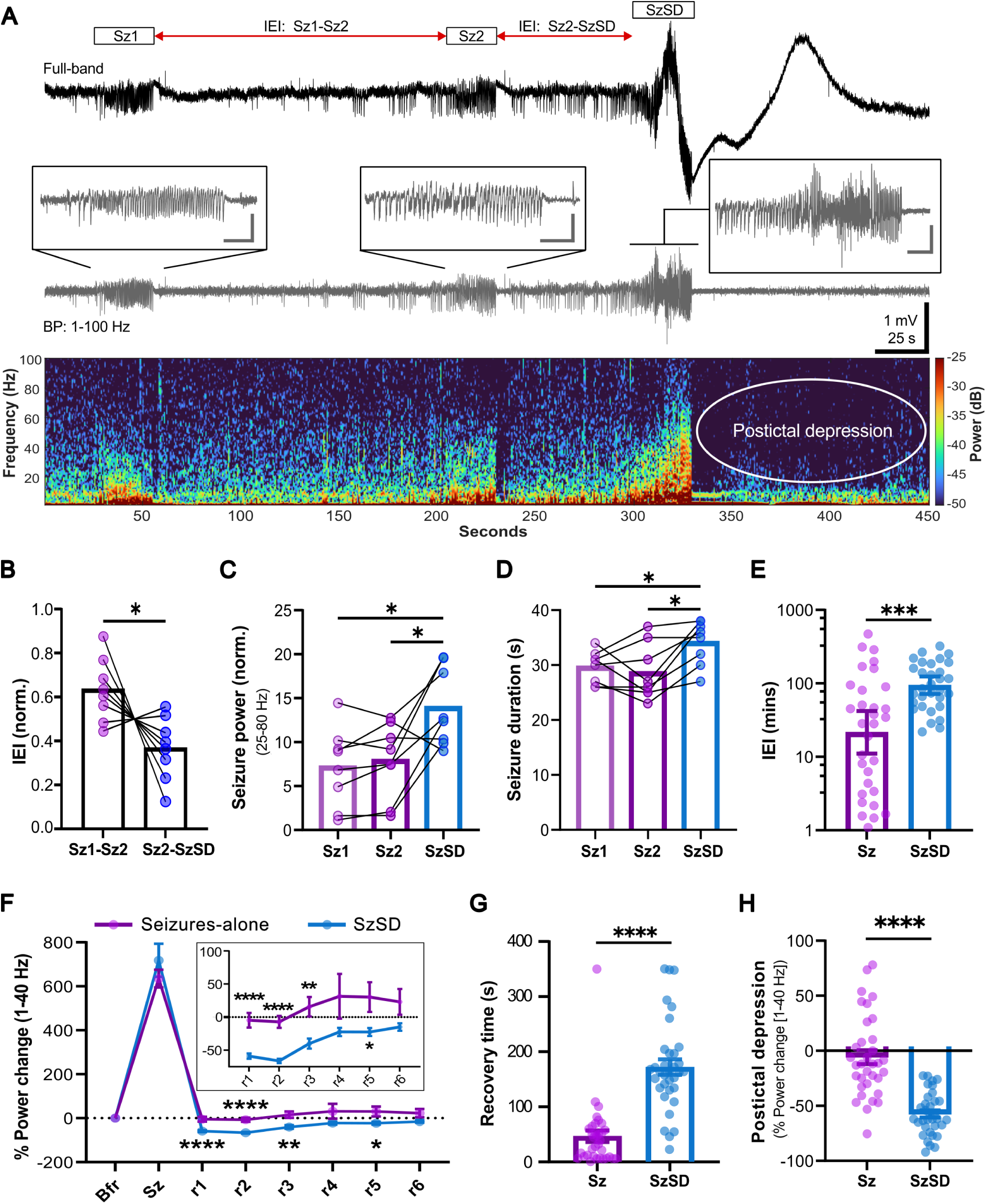
Spontaneous seizures associated with spreading depolarization exhibit increased power, pronounced postictal depression, and longer inter-event intervals. (**A**) Representative full-band iEEG recording showing two seizure-alone events (Sz1 and Sz2) preceding a seizure associated with spreading depolarization (SzSD). *Top*: full-band trace illustrating progressively shorter inter-event intervals (IEIs) before SzSD. *Middle*: corresponding 1-100 Hz filtered trace with expanded views of individual seizures. *Bottom*: corresponding spectrogram showing increased seizure power during SzSD and pronounced postictal depression. (**B**) IEIs progressively shortened before SzSD events. Normalised IEI: Sz1-Sz2, 0.63±0.05; Sz2-SzSD, 0.37±0.05; Paired t-test, \**P* = 0.0338, *n =* 8. (**C**) SzSD events exhibited greater seizure power (25-80 Hz) than preceding seizure-alone events. Seizure power (25-80 Hz, normalised): Sz1, 7.18±1.59; Sz2, 7.93±1.50; SzSD, 13.96±1.57; RM one-way ANOVA: *F*(1.13, 7.92) = 12.12, *P* = 0.0074; *n* = 8; Tukey’s multiple comparisons: Sz1 vs Sz2, *P* = 0.4574; Sz1 vs SzSD, \**P* = 0.0115; Sz2 vs SzSD, \**P* = 0.0390. (**D**) SzSD events exhibited longer seizure duration than preceding seizure-alone events. Seizure duration: Sz1, 29.63±1.05 s; Sz2, 28.63±1.81 s; SzSD, 34.13±1.43 s; RM one-way ANOVA: *F*(1.99, 13.94) = 7.25, *P* = 0.0070; *n* = 8; Tukey’s multiple comparisons: Sz1 vs Sz2, *P* = 0.8027; Sz1 vs SzSD, \**P* = 0.0449; Sz2 vs SzSD, \**P* = 0.0237. (**E**) Inter-event intervals following SzSD events were significantly longer than those following seizure-alone (Sz) events. Sz: 75.42±20.66 min, *n* = 31; SzSD: 120.80±14.98 min, *n* = 31. Mann-Whitney test, \*\*\**P* = 0.0008. Bars and error lines represent geometric mean and 95% confidence intervals (geometric mean [lower 95%CI, upper 95%CI]): Sz, 21.56 [11.08, 41.95] mins; SzSD, 94.17 [71.62, 123.80] mins. (**F**) Postictal electrographic recovery was prolonged following SzSD events. Percentage power change before (Bfr), during (Sz), and after seizures (r1-r6) revealed greater and more persistent postictal suppression after SzSD. Linear mixed-effects analysis: *F*(1.31, 81.90) = 215.4, *P* <0.0001; Tukey’s multiple comparisons, Sz vs SzSD: r1 and r2, \*\*\*\**P* <0.0001; r3, \*\**P* = 0.0014; r5, \**P* = 0.0307. For mean±SEM and all comparisons, see Supplementary Table 6. (**G**) Recovery time of postictal electrographic activity was significantly prolonged following SzSD events. Recovery time: Sz, 45.69±10.26 s, *n* = 35; SzSD, 173.50±13.64 s, *n* = 35. Mann-Whitney test, \*\*\*\**P* < 0.0001. (**H**) SzSD events exhibited markedly greater postictal depression (PID):. Sz, -4.37±5.72%, *n* = 35; SzSD, -58.16±3.33%, *n* = 35; Mann-Whitney test, \*\*\*\**P* <0.0001.

We next investigated whether seizures associated with SDs exhibit distinct electrophysiological signatures compared with seizure-alone events, and whether these features are conserved across hippocampal and cortical recording sites. Across both regions, seizures associated with SDs displayed significantly longer duration and greater coastline measures compared with seizures occurring in isolation (Fig. 7A; Supplementary Fig. 7; Supplementary Table 7). In contrast, seizure-associated spectral power differed between recording regions (Supplementary Fig. 7; Supplementary Tables 8-11). Cortical recordings showed a significant increase in spectral power during SD-associated seizures, whereas hippocampal recordings exhibited no additional increase (Fig. 7B; Supplementary Table 12). Notably, hippocampal seizure-alone events already displayed power levels comparable to those observed during cortical SD-associated seizures. Despite these differences in seizure power, postictal depression was markedly greater following seizure-associated SDs in both hippocampal and cortical recordings (Fig. 7C; Supplementary Table 13).

**Figure 7.**
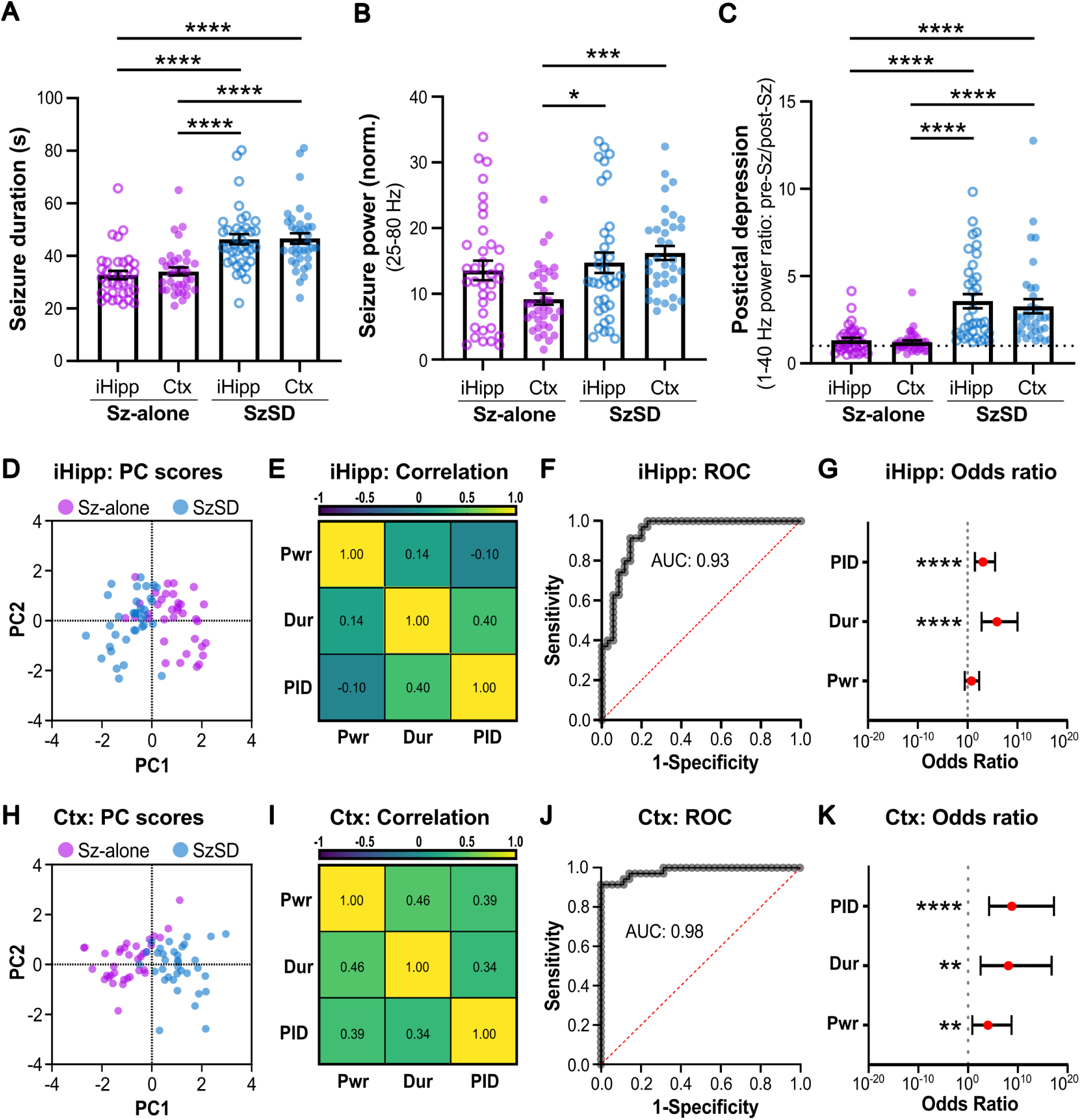
Principal component and multivariable analyses reveal distinct electrographic signatures that distinguish spontaneous seizure types in hippocampal and cortical regions of Kir4.1-cKO mice. (**A**) Seizures associated with spreading depolarization (SzSD) exhibited significantly longer seizure duration than seizure-alone events in both ipsilateral hippocampal (iHipp) and cortical (Ctx) recordings. Two-way ANOVA, *F*(1, 138) = 53.21; *P* < 0.0001. Tukey’s multiple comparisons test, \*\*\*\**P* <0.0001. For mean±SEM and all multiple comparisons, see Supplementary Table 7. (**B**) Cortical SzSD events exhibited greater seizure power than seizure-alone events, whereas seizure power did not differ between seizure types in the ipsilateral hippocampus. Two-way ANOVA, *F*(1, 138) = 10.15; *P* = 0.0018. Tukey’s multiple comparisons test: \**P* = 0.0198, \*\*\**P* = 0.0006. For mean±SEM and all multiple comparisons, see Supplementary Table 12. (**C**) SzSD events exhibited significantly greater postictal depression (PID) than seizure-alone events in both hippocampal and cortical recordings. Two-way ANOVA, *F*(1, 136) = 51.64; *P* <0.0001. Tukey’s multiple comparisons test: \*\*\*\**P* <0.0001. For mean±SEM and all comparisons, see Supplementary Table 13. (**D**) Principal component analysis (PCA) of hippocampal seizure events (*n* = 70) using seizure power, seizure duration, and PID revealed distinct clustering of seizure-alone (*n* = 35) and SzSD (*n* = 35) events. PC1 and PC2 explained 45.80% and 36.03% of total variance, respectively (Supplementary Fig. 8). (**E**) Spearman’s rank correlation matrix for hippocampal seizure events show a positive correlation between seizure duration (Dur) and PID (*P* = 0.005), whereas seizure power (Pwr) was weakly correlated with both variables (*n* = 70 events; For all *P-*values, see Supplementary Table 14). (**F**) Receiver operating characteristic (ROC) show strong discrimination between seizure-alone and SzSD events in the hippocampal region (AUC = 0.93, *P* < 0.0001, *n* = 70 events). (**G**) Odds ratio analysis of the hippocampal logistic regression model demonstrated that greater PID and longer seizure duration significantly increased the odds of classification as an SD-associated seizure (PID, \*\*\*\**P* < 0.0001; seizure duration, \*\*\*\**P* < 0.0001; seizure power, *P* = 0.2751; *n* = 70 events). For odds ratios and 95% confidence intervals, see Supplementary Table 16. (**H**) PCA of cortical seizure events (*n* = 70), using seizure power, seizure duration and PID, revealed distinct clustering of seizure-alone (*n* = 35) and SzSD (*n* = 35) events. PC1 and PC2 explained 57.18% and 25.93% of the total variance, respectively (Supplementary Fig. 8). (**I**) Spearman’s rank correlation matrix for cortical seizure events demonstrate positive correlations between seizure power (Pwr), seizure duration (Dur) and PID. *n =* 70 events. For all *P-*values, see Supplementary Table 15. (**J**) ROC show strong discrimination between seizure-alone and SzSD events in the cortical region (AUC = 0.98, *P* < 0.0001, *n* = 70 events). (**K**) Odds ratio analysis of the cortical logistic regression model demonstrated that greater PID, longer seizure duration, and higher seizure power were each significantly associated with increased odds of classification as an SD-associated seizure (PID, \*\*\*\**P* <0.0001; seizure duration, \*\**P* = 0.0017; seizure power, \*\**P* = 0.0074. For odds ratios and 95% confidence intervals, see Supplementary Table 16.

We next investigated whether electrophysiological features measurable using conventional AC-coupled recordings could distinguish seizures associated with SDs from seizures-alone events, using DC-coupled recordings as the ground-truth indicator of SD occurrence. Principal component analysis incorporating seizure power, seizure duration, and postictal depression demonstrated clear separation between seizure-alone and SD-associated seizure events in both hippocampal and cortical recordings, indicating distinct electrophysiological signatures associated with the two types of seizure events (Fig. 7D, H; Supplementary Fig. 8). Correlation analyses revealed region-specific relationships between these electrophysiological features. In hippocampal recordings, seizure duration positively correlated with postictal depression, whereas cortical recordings demonstrated strong positive correlations between seizure power, seizure duration, and postictal depression (Fig. 7E, I; Supplementary Table 14, 15).

To determine whether these electrophysiological features could retrospectively identify SD-associated seizures, we performed multivariable logistic regression analyses using seizure power, seizure duration, and postictal depression as predictor variables. These models discriminated seizures associated with SDs from seizure-alone events in both hippocampal and cortical recordings (ROC: iHipp, AUC = 0.93, *P* < 0.0001; Ctx, AUC = 0.98, *P* < 0.0001; Fig. 7F, J; Supplementary Fig. 9). Odds ratio estimates indicated that, in hippocampal recordings, longer seizure duration and greater postictal depression increased the likelihood of classification as an SD-associated seizure. In cortical recordings, increased seizure duration, seizure power, and postictal depression were each associated with classification as an SD-associated seizure (Fig. 7G, K; Supplementary Table 16).

Together, these findings demonstrate that SD-associated seizures exhibit distinct electrophysiological signatures that are detectable using conventional AC-coupled recordings, supporting robust discrimination of SD-associated seizures and seizure-alone events despite the absence of DC measurements.

### Postictal behavioural impairments after spontaneous seizures with spreading depolarizations

Using time-locked video recordings acquired during continuous DC-coupled telemetry, we next investigated behavioural correlates of seizures occurring with and without spreading depolarizations (SDs). Behavioural analysis was performed on seizure-alone and seizure-associated SD events (12 events per group), with behaviour assessed both during seizures and throughout the first 120 seconds following electrographic seizure termination (postictal period). Seizure severity was quantified using a modified Racine scale, while postictal behaviour was categorised as exploratory behaviour, behavioural arrest, or abnormal behaviour (see Methods).

Seizure and postictal behavioural profiles differed markedly between seizure-alone and SD-associated seizure events (Fig. 8A). SD-associated seizures exhibited substantially greater behavioural severity, reaching significantly higher maximum Racine scores compared with seizure-alone events (Fig. 8B). Consistent with this, seizure-alone events spent the majority of seizure time in lower non-convulsive Racine stages (stages 1-2), whereas SD-associated seizures spent a markedly greater proportion of seizure time in higher convulsive stages (stages 3-6; Fig. 8C; Supplementary Fig. 10; Supplementary Table 17). Together, these findings demonstrate that SD-associated seizures are linked to a substantially more severe convulsive seizure phenotype.

**Figure 8.**
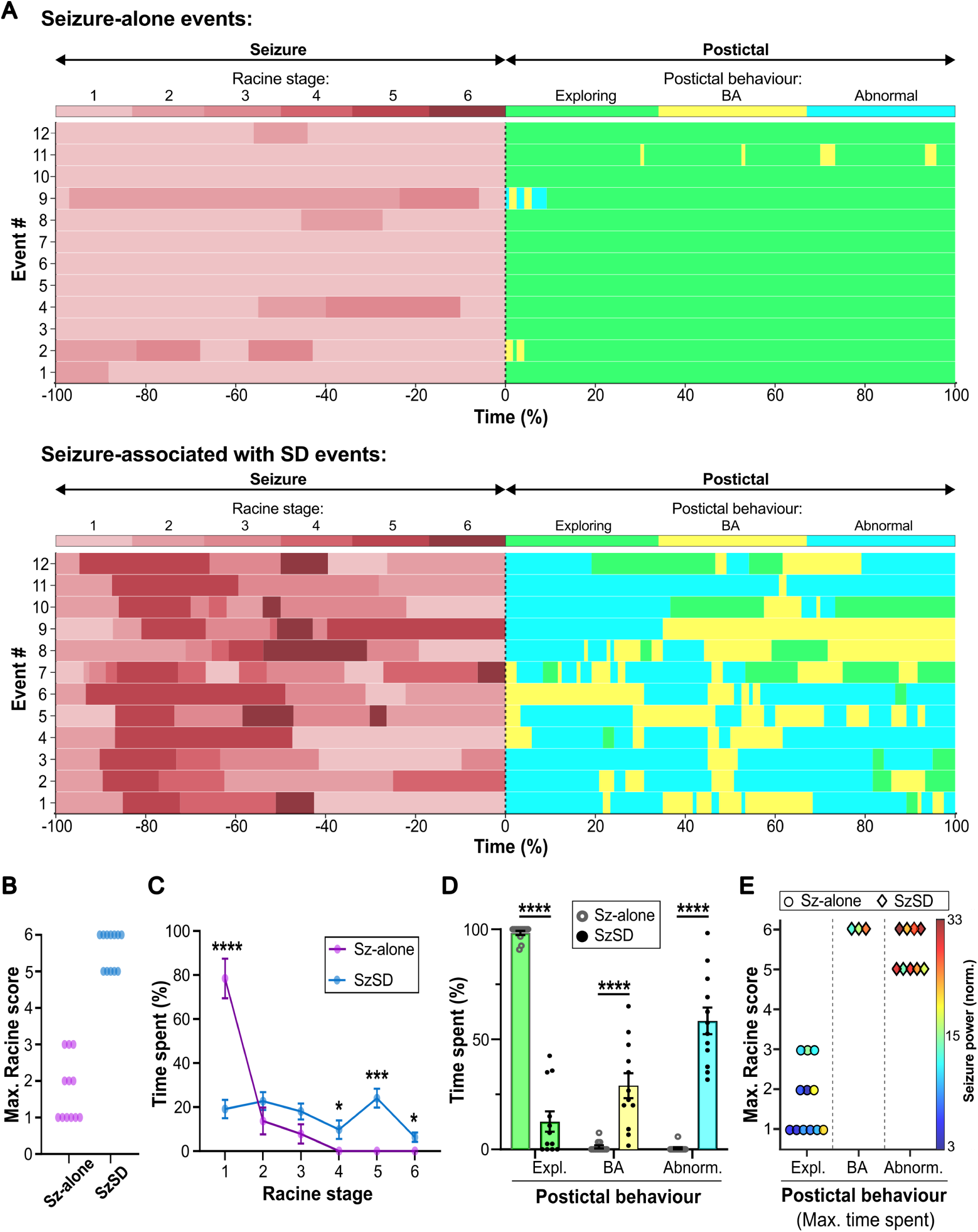
Severe convulsive seizures associated with spreading depolarization exhibit marked postictal behavioural impairment. (**A**) Heatmaps showing seizure Racine stage during the course of the seizure (time: -100 to 0 %) and postictal behaviour during the 120-second period following seizure termination (time: 0-100%). Time 0% denotes seizure end. Seizure-alone, *n* = 12 events; SzSD, *n* = 12 events (6 mice/group). (**B**) SzSD events exhibited a more severe convulsive phenotype, with higher maximum Racine scores than seizure-alone events. Perecntage of events in Racine scores (RS); Sz: RS1=50%, RS2=25%, RS3=25%; SzSD: RS5=42%, RS6=58%; Chi-square test, \*\*\*\**P* <0.0001. (**C**) Mice showing SzSD spent significantly greater proportion of seizure time in higher Racine stages (stages: 4-6), whereas seizure-alone events were predominantly associated with Racine stage-1. Two-way ANOVA, *F*(3.43, 75.39) = 9.49; *P* < 0.0001; Tukey’s multiple comparisons, Sz-alone vs SzSD: Racine stage-1, \*\*\*\**P* <0.001; Racine stage-4, \**P* = 0.0407; Racine stage-5, \*\*\**P* = 0.0002; Racine stage-6, \**P* = 0.0123. For mean±SEM and all comparisons, see Supplementary Table 17. (**D**) Postictal behaviour was markedly altered following SzSD events, with mice spending most of the 120-s postictal period displaying abnormal behaviour or behavioural arrest, whereas seizure-alone events were associated predominantly with exploratory behaviour. Two-way ANOVA, *F* (2, 66) = 54.87, *P* < 0.0001; Tukey’s multiple comparisons, Sz-alone vs SzSD: exploratory behaviour, \*\*\*\**P* < 0.0001, BA, \*\*\*\**P* < 0.001; abnormal behaviour, \*\*\*\**P* < 0.0001. For all mean±SEM and comparisons, see Supplementary Table 18. (**E**) Postictal behavioural severity was significantly associated with maximum seizure Racine score and seizure power (ordinal logistic regression: χ^2^(2) = 29.2; *P* <0.001, *n =* 24 events). Seizure type (Sz-alone vs SzSD) showed complete segregation with postictal behavioural severity (Fisher’s exact test, *P* <0.0001; *n =* 24 events). See Supplementary Fig. 11 for ordinal logistic regression analyses showing the relationship between maximum seizure Racine score, seizure-associated spectral power, and postictal behavioural severity.

Postictal behaviour was assessed during the first 120 seconds following electrographic seizure termination. Following seizure-alone events, mice rapidly resumed exploratory behaviour and spent almost the entire postictal period actively exploring (98.33±0.95%; Fig. 8A, D; Supplementary Table 18; Supplementary movie S1). In contrast, seizures associated with SDs were followed by profound behavioural impairment, with mice spending the majority of the postictal period either behaviourally arrested or displaying abnormal behaviours (Fig. 8D). Thus, seizure-associated SDs consistently represented the most severe seizure and post-ictal behavioural phenotype observed in Kir4.1-cKO mice (see Supplementary movies S2-5).

To determine whether postictal impairment was associated with seizure severity, ordinal logistic regression analysis was performed using maximum Racine score and seizure power as predictors of postictal behavioural severity. Maximum Racine score and seizure power significantly predicted postictal behavioural impairment, with increasing seizure severity associated with more pronounced postictal deficits (χ²(2) = 29.2, *P* < 0.001, *n* = 24 events; Fig. 8E; Supplementary Fig. 11). Consistent with these findings, seizure-alone and SD-associated seizure events showed complete separation based on postictal behavioural severity (Fig. 8E; Fisher’s exact test, *P* < 0.0001), with SD-associated seizures producing the most severe postictal behavioural impairments.

Collectively, these findings demonstrate that focal astrocytic Kir4.1 loss disrupts potassium homeostasis, promotes epileptic hyperexcitability, and increases susceptibility to spreading depolarizations. Seizures associated with spreading depolarizations were characterised by increased seizure duration, enhanced cortical seizure power, more severe convulsive behavioural phenotypes, prolonged postictal depression, and profound postictal behavioural impairments.

## Discussion

Astrocytic dysfunction is increasingly recognised as a critical contributor to epileptogenesis and seizure dynamics,^1^ yet the extent to which focal astrocytic pathology alone is sufficient to drive spontaneous seizures in the mature brain has remained unclear. Here, we demonstrate that focal loss of astrocytic Kir4.1 within the adult hippocampus disrupts extracellular potassium buffering, induces spontaneous recurrent seizures, and promotes seizure-associated spreading depolarizations (SDs). Importantly, SD-associated seizures were characterised by greater behavioural seizure severity, profound postictal depression, and postictal behavioural impairment. Together, these findings suggest that more severe seizures are more likely to be associated with transitions into SD states, which are accompanied by markedly worsened postictal dysfunction. These results establish astrocytic potassium buffering as a critical regulator of seizure susceptibility and further support an emerging role for Kir4.1-mediated potassium homeostasis in regulating SD vulnerability and pathological postictal network suppression. Consistent with this, recent work by Tyurikova *et. al.*^6^ demonstrated that astrocytic Kir4.1 expression critically regulates extracellular potassium dynamics, excitatory synaptic transmission, and SD propagation, with Kir4.1 overexpression markedly attenuating high-potassium SD waves and activity-dependent network excitability. Together, these studies support a model in which astrocytic Kir4.1 dysfunction destabilises activity-dependent ionic homeostasis, promoting both seizure generation and seizure-SD coupling.

### Astrocytic Kir4.1 dysfunction impairs potassium buffering and promotes epileptogenesis

Kir4.1 channels are highly enriched within astrocytic perisynaptic and perivascular processes, where they play a central role in spatial potassium buffering and maintenance of extracellular ionic homeostasis.^2,3^ In addition to regulating extracellular potassium levels, Kir4.1 channels help maintain the hyperpolarised astrocytic membrane potential required for efficient glutamate uptake, thereby coupling potassium and glutamate homeostasis within the tripartite synapse.^30^ Reduced astrocytic Kir4.1 expression and impaired Kir-mediated currents have been consistently identified in human temporal lobe epilepsy and hippocampal sclerosis, supporting a mechanistic role for astrocytic potassium buffering dysfunction in epilepsy.^2,8^

In the present study, we used focal conditional deletion of Kir4.1 within the adult hippocampus to avoid the widespread developmental abnormalities associated with constitutive knockout models,^4^ and instead isolate the consequences of acquired astrocytic dysfunction within established neural circuits. Importantly, Kir4.1 reduction in the present model was partial rather than complete, producing an approximately 50% reduction in protein expression comparable to that reported in human temporal lobe epilepsy tissue.^7^ Mechanistically, Kir4.1 loss substantially impaired activity-dependent extracellular potassium buffering, resulting in larger and more prolonged extracellular potassium transients following synaptic stimulation. These findings are consistent with prior work demonstrating that Kir4.1 critically regulates astrocytic potassium uptake and local excitatory transmission.^6^ Elevated extracellular potassium is predicted to depolarise neuronal membranes, facilitate recurrent excitation, and lower seizure threshold.^2^ Consistent with this, Kir4.1-cKO mice exhibited increased susceptibility to evoked seizures and developed spontaneous recurrent seizures *in vivo*. Our findings therefore support the concept that focal astrocytic dysfunction alone can be sufficient to drive spontaneous epileptic activity in the adult brain, highlighting astrocytes not simply as modulators of neuronal excitability but as direct contributors to epileptogenesis.^1^

### Context-dependent relationships between seizures and spreading depolarizations

Although previous studies have reported that seizures associated with SDs are shorter in duration and proposed that SDs may represent an endogenous anti-seizure mechanism capable of terminating ictal activity, these findings were derived primarily from acute chemoconvulsant models in otherwise healthy brain tissue.^16,17^ One possible explanation is that SD initiation thresholds may differ substantially between relatively naïve and chronically epileptic tissue. Although epileptic brain networks are highly seizure-prone, previous *in vitro* studies of rodent and human epileptic tissue have demonstrated increased potassium thresholds for SD initiation despite enhanced seizure susceptibility.^31-34^ In relatively naïve tissue, lower SD thresholds may permit earlier SD recruitment during seizure evolution, potentially curtailing seizure duration. In our optogenetic experiments in Kir4.1-cKO mice, we similarly observed a trend towards shorter seizure duration when SDs were present, although this did not reach statistical significance. Importantly, in these experiments SD emerged within the same hippocampal network in which seizures were initiated (Supplementary Fig. 12), supporting the hypothesis that locally generated SDs may contribute directly to seizure termination or suppression. In contrast, spontaneous seizure recordings in chronically epileptic animals revealed the opposite relationship. Here, seizures associated with SDs were significantly longer than seizure-alone events. Notably, SDs were never first detected in the hippocampus, the presumed seizure onset zone, but were consistently first observed in the cortex and occurred during more severe seizures. These findings suggest that in the chronically epileptic brain, seizure-associated SDs may not simply represent a locally acting seizure-terminating mechanism, but instead may emerge secondary to prolonged or more intense seizure propagation into distributed cortical networks. A higher SD initiation threshold in chronically epileptic tissue may further contribute to this phenomenon, requiring greater seizure intensity, duration, or spatial recruitment before SD generation occurs. Consistent with this, SD-associated seizures during acutely evoked optogenetic events were shorter in duration than in SD-associated spontaneous seizures (Supplementary Fig. 12B). Together, these observations indicate that the relationship between seizures and SDs is likely highly dependent on both the underlying disease state and the spatial relationship between seizure onset and SD initiation. More broadly, these findings support the concept that seizure susceptibility and SD susceptibility may represent mechanistically dissociable properties of epileptic networks.

### Spreading depolarizations as seizure cluster-terminating events

The observation that recurrent seizure clusters culminated in SD-associated seizures is particularly notable. Seizure clustering can occur in both chronic animal models and human epilepsy. Prior work in a rat model of temporal lobe epilepsy showed that cluster-terminating seizures are associated with greater behavioural severity, increased cortical EEG power, and broader network recruitment, although AC-coupled recordings could not determine whether these terminal seizures were associated with SDs.^35^ Our findings suggest that SD may be one mechanism contributing to this cluster-terminating seizure phenotype. This interpretation is consistent with preclinical studies showing that seizures terminating with SD are followed by a prolonged refractory period before subsequent seizures, supporting the concept of SD as an endogenous anti-seizure mechanism that suppresses pathological network activity and moves the brain away from a seizure-permissive state.^16,17^ In our chronic spontaneous epilepsy model, SD-associated seizures were similarly followed by longer intervals before the next seizure, extending this principle beyond acute chemoconvulsant and *ex vivo* models. However, this apparent network-stabilising effect came at a substantial physiological cost, with SD-associated seizures producing profound postictal suppression and behavioural impairment. This may be clinically important because seizure clustering has been linked to SUDEP risk in both animal models and human studies, although this relationship is likely context-dependent rather than simply causal.^36-38^ Consistent with this cautious interpretation, we observed one SUDEP event during DC-coupled telemetry in which the fatal seizure occurred as an SD-associated cluster-terminating event (see Supplementary Fig. 13); however, given the observational nature of this finding, no mechanistic conclusions regarding SUDEP can be drawn.

### Postictal impairment and SUDEP risk

SD-associated seizures were associated with substantially more severe postictal dysfunction both electrophysiologically and behaviourally, including prolonged depression of electrographic activity, delayed recovery of baseline network dynamics, impaired postictal behaviour, and delayed recovery. These findings are consistent with reports from both rodent and human studies linking seizure-associated SDs to prolonged postictal suppression, impaired arousal, abnormal ambulation, and delayed behavioural recovery,^22^ and extend these observations by demonstrating that SD occurrence robustly segregates severe versus mild postictal phenotypes in a chronic spontaneous epilepsy model. However, SD is unlikely to be the sole mechanism underlying postictal dysfunction. Postictal depression can occur in the absence of SD and has been linked to multiple complementary processes, including seizure-induced adenosine accumulation, dysregulated inhibitory neurotransmission, metabolic stress, and neurovascular disturbances.^39-42^ Furthermore, because seizure severity is likely to influence both SD recruitment and other candidate mediators of postictal dysfunction, the present data do not permit determination of the relative contribution of SD to the severe impairment observed following SD-associated seizures. Nevertheless, recent experimental studies have shown that seizure-associated SD can directly drive postictal behavioural abnormalities, supporting a causal role for SD in at least some postictal manifestations.^22^ Together, these findings support a model in which SD contributes to postictal dysfunction, potentially acting in concert with other seizure-associated mechanisms to exacerbate postictal impairment and prolong recovery following particularly severe seizures.

The profound postictal suppression associated with SD-associated seizures may also have important implications for SUDEP. Increasing evidence from genetic SUDEP-prone mouse models implicates spreading depolarization in terminal brainstem dysfunction during lethal seizures, resulting in apnoea and seizure-associated failure of cardiorespiratory control.^18-21,43,44^ Although the present study was not designed to directly address SUDEP mechanisms, the marked postictal suppression and behavioural impairment associated with SD-coupled seizures in our model are consistent with this emerging framework. Future studies incorporating brainstem recordings and cardiorespiratory monitoring will be required to directly test these possibilities.

### Translational implications of SD-associated seizure signatures

An important translational finding of the present study is that electrophysiological features measurable using conventional AC-coupled recordings robustly discriminated SD-associated seizures from seizure-alone events, despite the absence of DC measurements. Using full-bandwidth DC-coupled recordings as ground-truth indicators of SD occurrence, we identified seizure duration, cortical seizure power, and postictal depression as key electrophysiological features associated with SD-associated seizures. If validated across additional preclinical models and human recordings, these signatures could enable retrospective identification of probable SD-associated seizures within existing AC-coupled electrophysiological datasets, substantially expanding understanding of SD prevalence and clinical significance without requiring widespread implementation of DC-coupled recording systems. The markedly enhanced postictal depression and behavioural impairment associated with SD-associated seizures further suggest that these events may represent a higher-risk seizure subtype associated with impaired arousal and greater postictal vulnerability. Importantly, convulsive seizures accompanied by postictal generalized EEG suppression (PGES) and postictal immobility have been associated with elevated SUDEP risk in clinical studies.^45,46^ While the postictal depression observed here should not be considered equivalent to PGES, given the limited spatial sampling and the persistence of residual electrophysiological activity, these findings raise the possibility that seizure-associated SDs may contribute to prolonged suppression of cortical activity and impaired behavioural recovery following seizures. Identification of electrophysiological biomarkers predictive of SD occurrence could therefore contribute to patient risk stratification and potentially guide therapeutic interventions targeting seizure-associated SD dynamics. More broadly, these findings suggest that spreading depolarizations may represent an under-recognised mechanistic driver of severe postictal depression and impaired behavioural recovery following seizures. Because SDs are largely invisible to conventional AC-coupled recordings, their contribution to postictal dysfunction and seizure severity may currently be substantially underestimated. Broader adoption of DC-coupled electrophysiological approaches, together with emerging graphene micro-transistor technologies capable of stable full-bandwidth recordings,^47^ may improve understanding of infraslow dynamics and seizure-SD coupling in human epilepsy.

### Technological advances to detect seizure-associated SDs

An important aspect of the present study is the use of full-bandwidth DC-coupled electrophysiology to investigate seizure-SD coupling *in vivo*. Conventional electrophysiological recordings frequently attenuate, distort or exclude infraslow activity through high-pass filtering, limiting detection of SDs and related infraslow dynamics.^48^ By using graphene-based micro-transistor arrays for acute DC-coupled recordings in awake head-fixed mice and chronic DC-coupled telemetry in freely moving animals, the present study enabled high-fidelity detection of seizures and SDs across complementary experimental paradigms. To enable long-term full-bandwidth recordings in freely moving animals, we employed a novel DC-coupled wireless telemetry platform capable of stable chronic recordings across multiple weeks. This approach overcomes important limitations of conventional tethered DC-recording systems, which are often restricted to shorter recording periods and may influence animal behaviour. The ability to perform continuous 24/7 DC-coupled recordings over extended periods was critical for capturing spontaneous SD-associated seizures and resolving their temporal, electrophysiological, and behavioural dynamics.

### Limitations

Several limitations should nevertheless be acknowledged. The spatial sampling of recording electrodes was limited, preventing precise determination of SD initiation sites and detailed mapping of its propagation dynamics. In addition, while the present study demonstrates robust seizure-SD coupling within a focal astrocytic Kir4.1-deficient model, further work will be required to determine whether similar electrophysiological signatures and postictal dynamics generalise across other chronic epilepsy models and human epilepsies. Extending these approaches to additional acquired and genetic epilepsy paradigms, together with human DC-coupled recordings, will be important for establishing the broader translational significance of SD-associated seizures.

In conclusion, the present findings demonstrate that focal astrocytic Kir4.1 dysfunction within the adult hippocampus is sufficient to induce spontaneous seizures, promote seizure-associated SDs, which profoundly worsen postictal recovery. These results position astrocytic potassium buffering as a critical regulator of pathological network-state transitions linking seizures, SDs, and postictal suppression. More broadly, they suggest that SD-associated seizures may represent a clinically important seizure subtype that remains substantially under-recognised using conventional electrophysiological approaches.

## Acknowledgements

We thank Prof. Matthew Walker (UCL) and Prof. Beate Diehl (UCL) for their valuable comments on the manuscript, and Dr. Olga Kopach (UCL) for help setting up protocols for *ex vivo* experiments.

## Supplementary figures

**Supplementary Figure 1.**
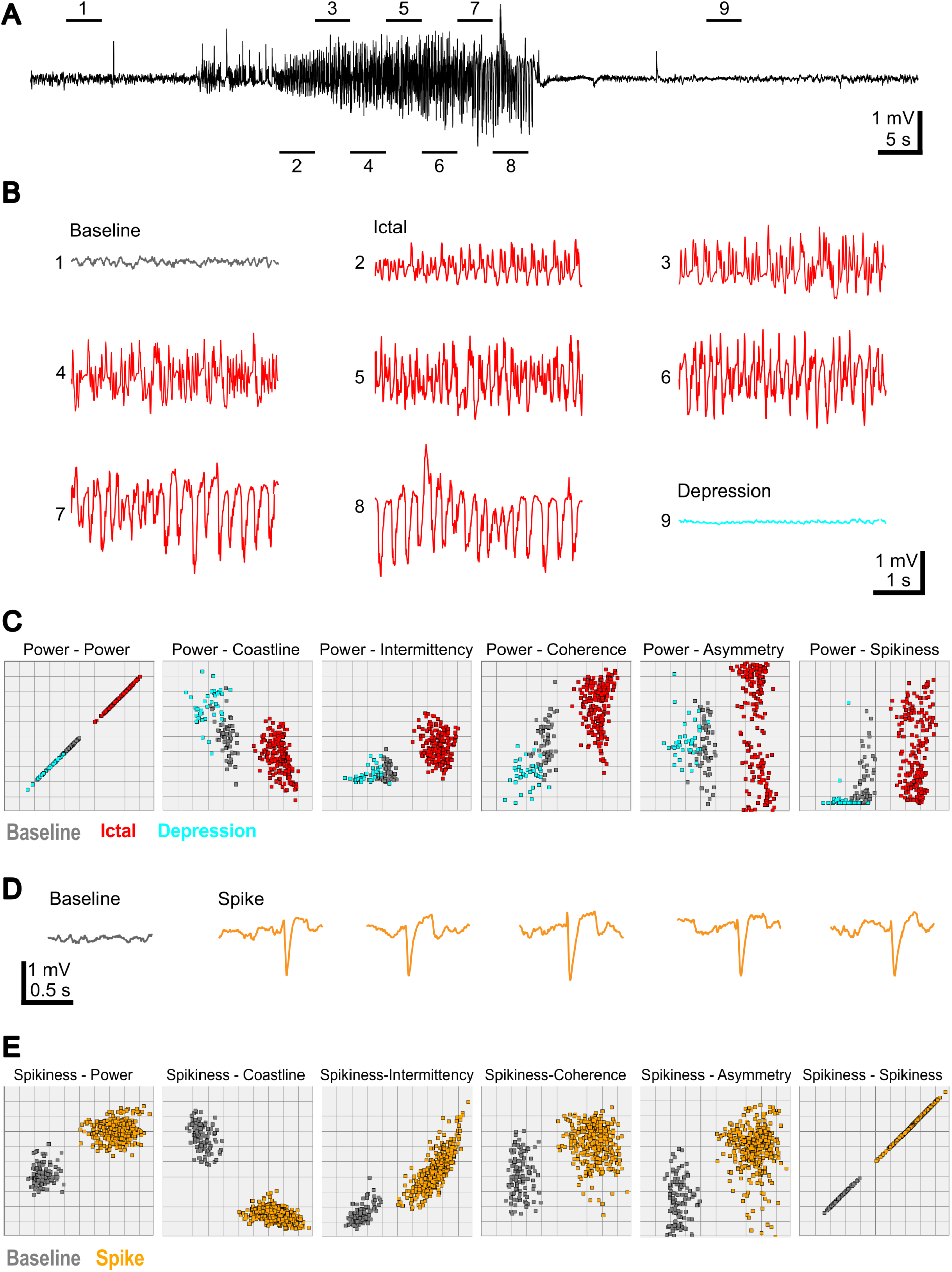
Automated analysis of seizures and spikes in Kir4.1-cKO mice using Neuroplayer Event Classifier tool in LWDAQ software (Open Source Instruments). (**A**) A representative seizure with 4-second window highlights (horizontal bars and numbers) of baseline, ictal and depression segments marked with numbers 1-9 for the seizure detection library. (**B**) Representative 4-second segments of the trace in **A** on a expanded timescale showing baseline (grey), ictal (red) and depression (cyan). (**C**) Final seizure library shown with 6 metrics to power: Power, coastline, intermittency, coherence, asymmetry and spikiness; with which semi-automated screening to select seizures from all iEEG data was performed. For more details on event classifier, see website: https://www.opensourceinstruments.com/Electronics/A3018/Neuroplayer.html#Event%20Classifier. (**D**) Representative 1-second baseline (grey) and spike (orange) segments for the spike detection library. (**E**) Final spike library shown with 6 metrics to spikiness: Power, coastline, intermittency, coherence, asymmetry and spikiness; with which automated screening to select spikes from all iEEG data was performed.

**Supplementary Figure 2.**
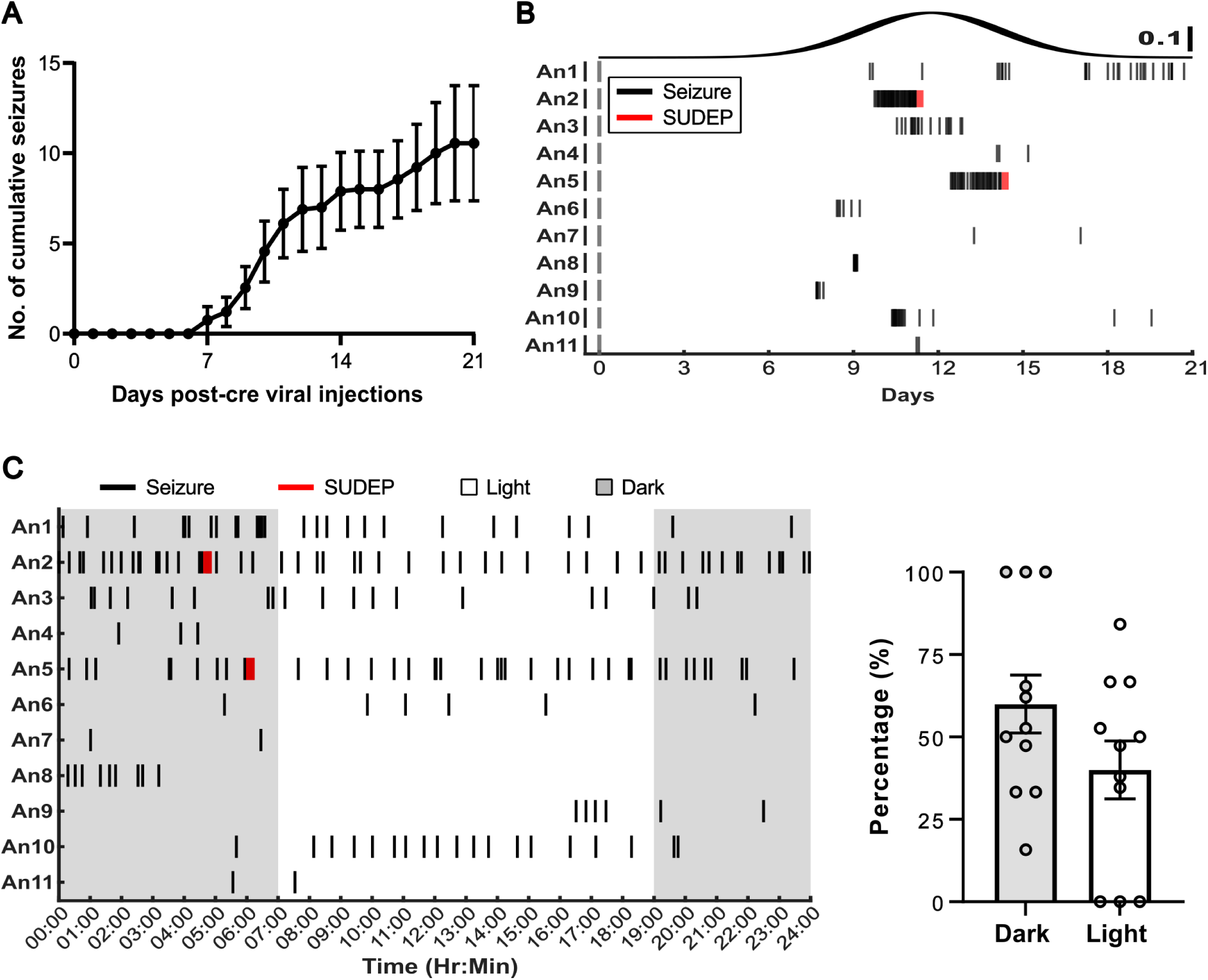
Evolution and circadian dynamics of spontaneous seizures recorded using AC-coupled subcutaneous transmitters for 21 days after Ast-Cre viral vector injection. (**A**) Cumulative seizure frequency plot show the development of spontaneous seizures around 7 days post-virus injection (dpi) in Kir4.1-cKO mice that continued to occur for the recording period (3 weeks/21 days). 21 dpi: cKO, 16.82±5.02 seizures, *n* = 11 mice. (**B**) Raster plot show the seizure development profile in individual animals and the probability density funtion plot show that seizures occurrence peaked at 11.76±2.42 dpi (*n* = 11 mice; value is represented as mean±SD). Given that seizure burden was maximal during the second post-injection week, subsequent DC-coupled recording experiments and quantitative analyses focused primarily on this time window. (**C**) *Left:* Circadian raster plot over 21 days show the occurrence of seizure events during light and dark phases. *Right*: Quantitative analysis of seizure event distribution reveal no significant difference in occurrence between light and dark phases. Dark, 59.99±8.80 %; Light, 40.01±8.80 %; paired t-test, *P* = 0.28; *n* = 11 mice.

**Supplementary Figure 3.**
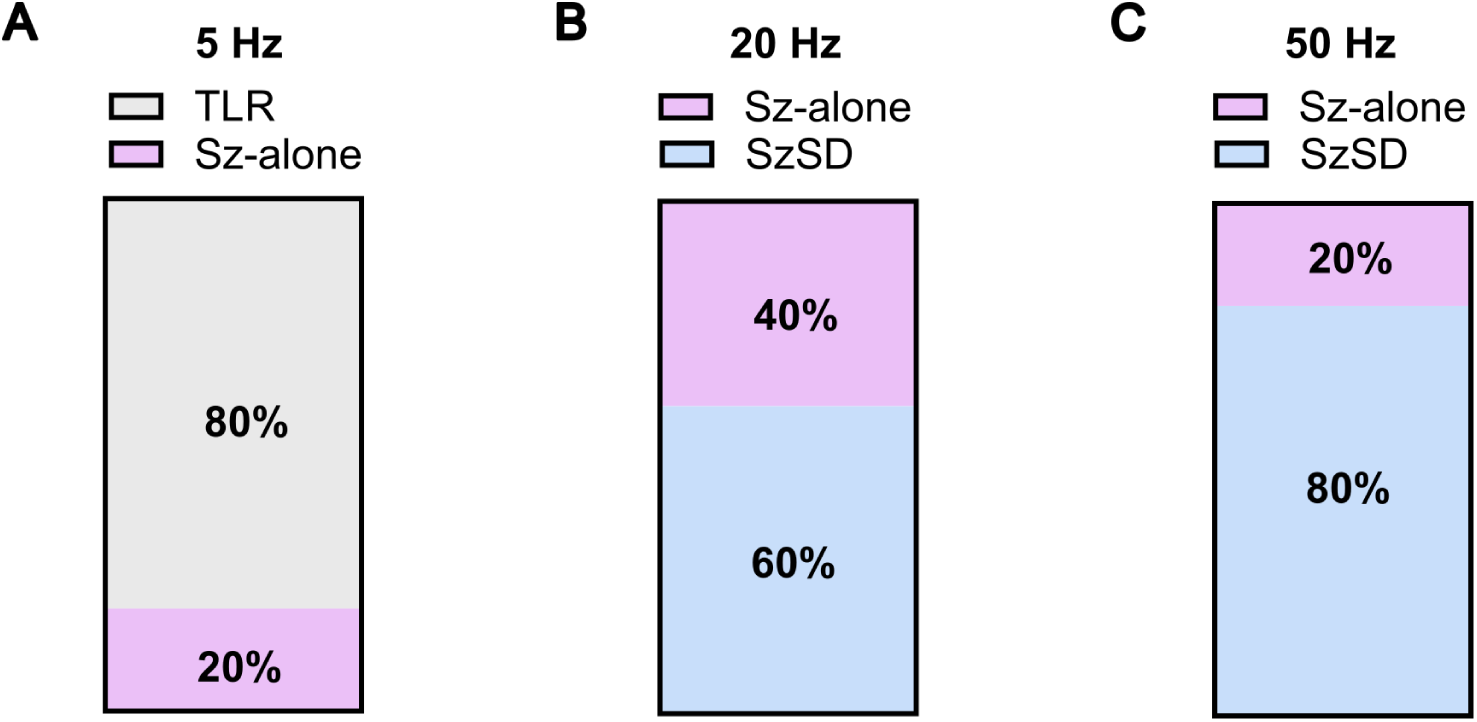
Optogenetic stimulation-evoked, frequency-dependent responses in awake, head-fixed Kir4.1-cKO mice. (**A**) 5 Hz-10 seconds stimulation (50 pulses; total energy, 1.37 mJ): 80% (4/5 trials) of the stimulation trials show only time-locked responses (TLR) whereas 20% (1/5 trials) induced a seizure-alone (Sz-alone) event. (**B**) 20 Hz-10 seconds stimulation (200 pulses; total energy, 5.46 mJ): All stimulation trials evoked seizure responses of which 60% (3/5 trials) are associated with spreading depolarization (SzSD) whereas 40% (2/5 trials) are seizure-alone events. (**C**) 50 Hz-10 seconds stimulation (500 pulses; total energy, 13.65 mJ): All stimulation trials evoked seizure responses of which 80% (4/5 trials) associated with spreading depolarization whereas 20% (1/5 trials) are seizure-alone events.

**Supplementary Figure 4.**
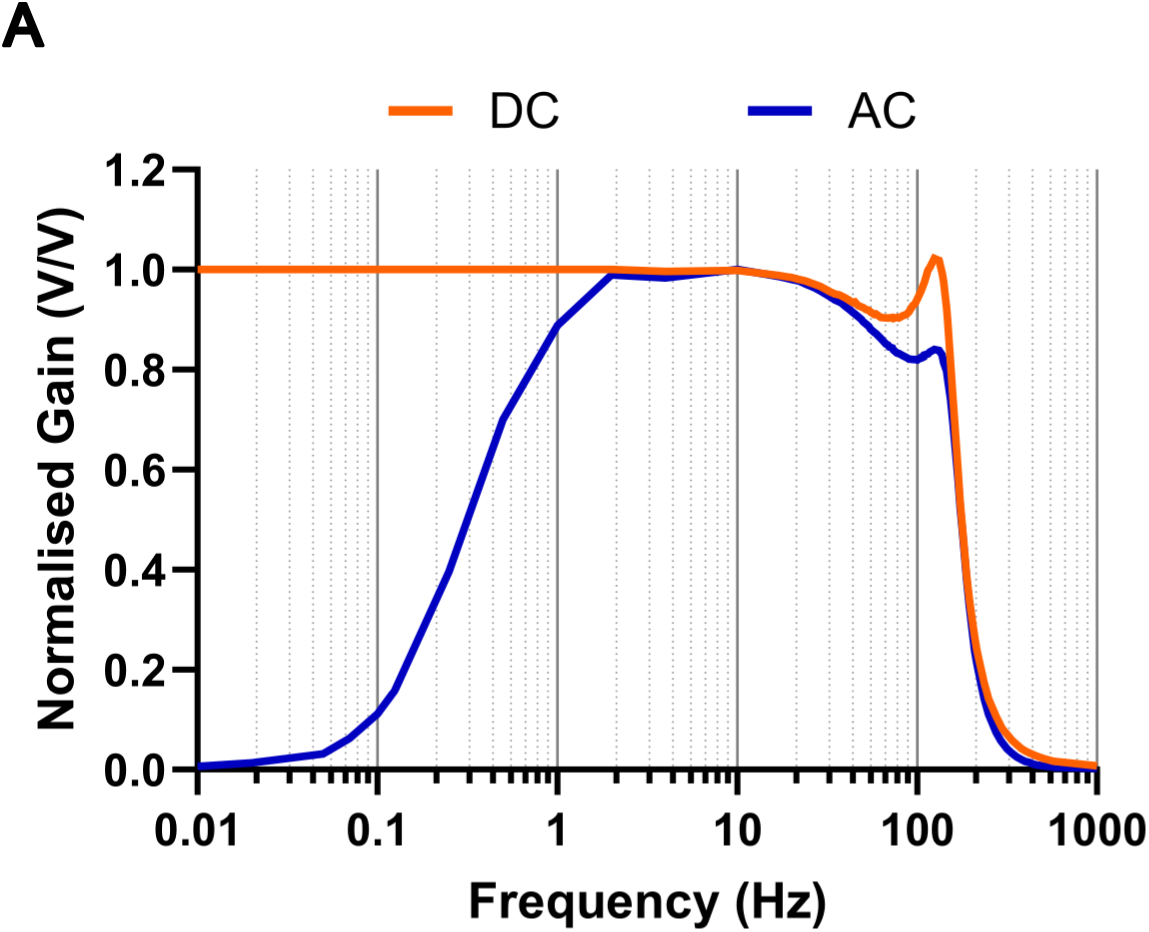
DC-coupled transmitters preserve low-frequency (<1 Hz) electrophysiological signals. (**A**) Frequency response curves of DC-coupled (orange) and AC-coupled (blue) telemetry transmitters, plotted as normalised voltage gain (V/V) across recording frequencies. DC-coupled transmitters maintained stable signal gain at low frequencies (<1 Hz), enabling recording of ultra-slow potential shifts such as spreading depolarizations. In contrast, AC-coupled transmitters showed marked attenuation of low-frequency signals below 1 Hz while preserving higher-frequency activity.

**Supplementary Figure 5.**
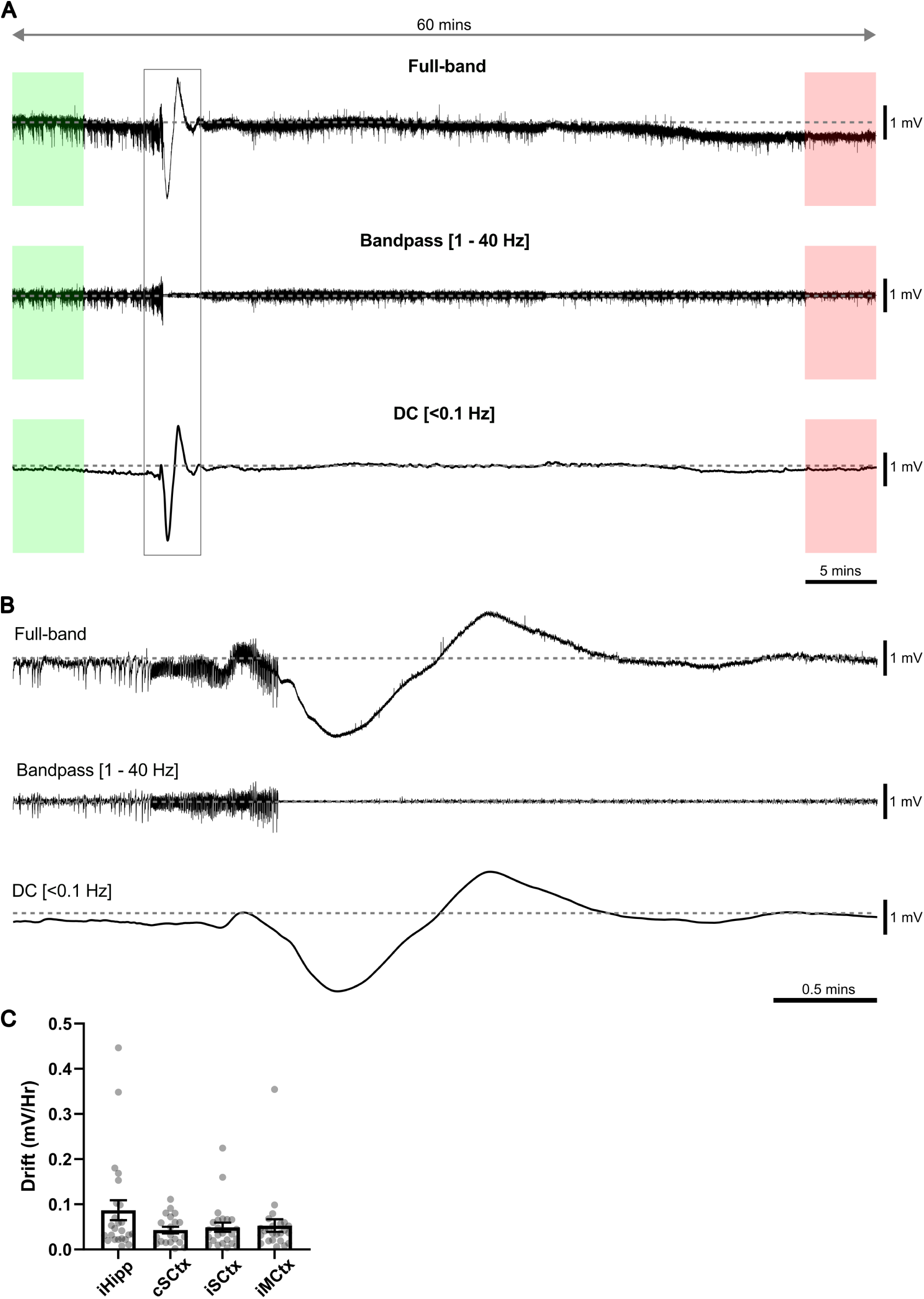
Stable chronic electrophysiological recordings with minimal signal drift using DC-coupled headmount transmitters. (A**) Representative one-hour** recording containing SD-associated seizure. *Top*, full-band raw trace; *middle*, bandpass-filtered (1-40 Hz); *bottom*, low-pass (DC) filtered (<0.1 Hz). Green and red shaded regions indicate the start and end windows used to calculate signal drift over the one-hour recording. (**B**) Expanded view of the electrophysiological signal highlighted (gray box) in panel A. (**C**) Signal drift (mV/hr) quantified across four recording electrodes in six mice over multiple recording days. Signal drift (‘*n’* values indicate the number of one-hour recordings): ipsilateral hippocampus, 0.09±0.02 mV/hr (*n* = 24); contralateral somatosensory cortex, 0.04±0.01 mV/hr (*n* = 20); ipsilateral somatosensory cortex, 0.05±0.01 mV/hr (*n* = 24); ipsilateral motor cortex, 0.05±0.01 mV/hr (*n* = 24). No significant difference across the four recording electrodes (Kruskal-Wallis statistic = 2.16, *P* = 0.54, total *n* = 92).

**Supplementary Figure 6.**
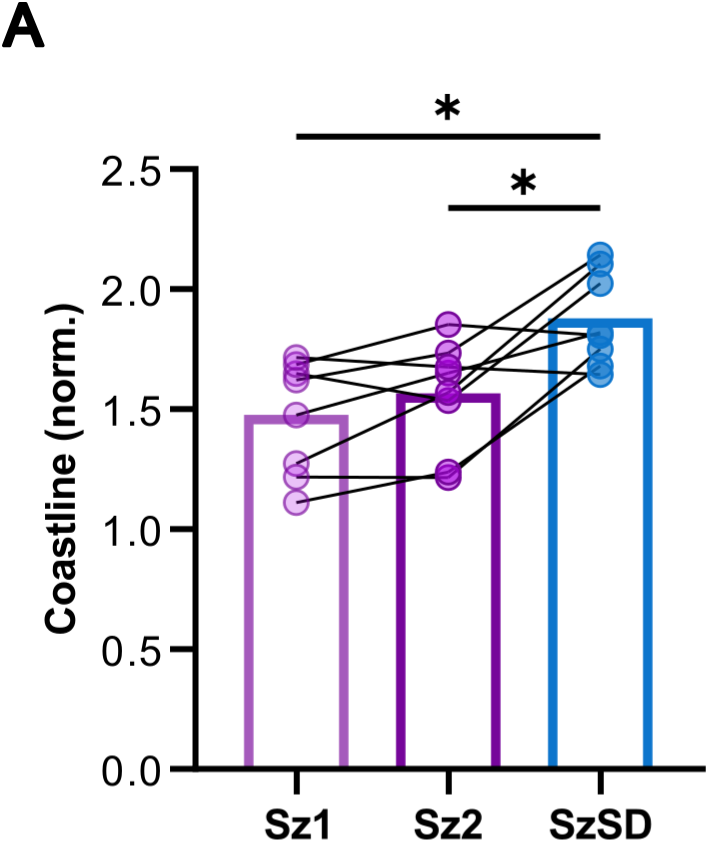
Spontaneous seizures associated with spreading depolarization (SzSD) are characterised by higher coastline. (**A**) Seizure associated with SD (SzSD) show higher coastline compared with the immediately preceding seizure-alone events (Sz1, Sz2). Event sequence: Sz1, Sz2, and SzSD. Coastline (norm.): Sz1, 1.47±0.08, *n* = 8; Sz2, 1.56±0.08, *n* = 8; SzSD, 1.87±0.07, *n* = 8; repeated measurements one-way ANOVA, *F* (1.357, 9.499) = 13.68, *P* = 0.0029; Tukey’s multiple comparisons test: Sz1 vs Sz2, *P* = 0.2048; Sz1 vs SzSD, \**P* = 0.0115; Sz2 vs SzSD, \**P* = 0.0211.

**Supplementary Figure 7.**
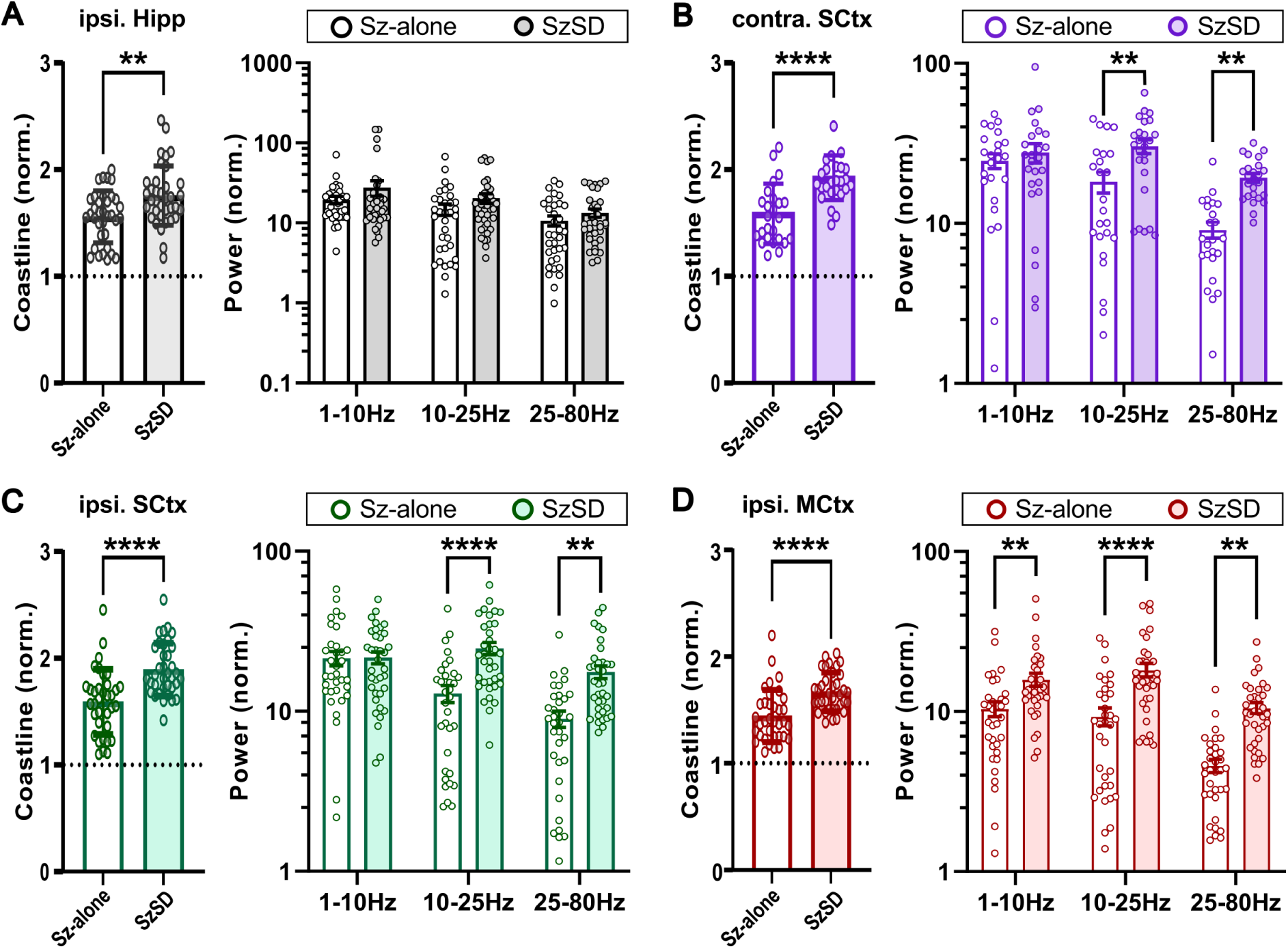
Spontaneous seizures with spreading depolarization detected in cortical regions show higher power compared with seizures-alone events. (**A**) SD-associated seizures (SzSD) recorded in ipsilateral hippocampus (ipsi. Hipp) show higher coastline (A, *left*) but similar power (A, *right*) compared to seizure-alone (Sz-alone) events. *Left*: Coastline (norm.): Sz-alone, 1.56±0.04, *n* = 35; SzSD: 1.76±0.05, *n* = 36; unpaired Welch’s t-test, \*\**P* = 0.02. *Right:* Power (norm.): Two-way ANOVA show main effects of frequency band (*F* (2, 204) = 6.671; *P* = 0.0016) and event type (F (1, 204) = 4.497; *P* = 0.0352). For mean±SEM and multiple comparisons, see Supplementary Table 8. (**B**) SzSD recorded in contralateral somatosensory cortex (contra. SCtx) show higher coastline (B, *left*) and power (B, *right*) compared to seizure-alone events. *Left:* Coastline (norm.): Sz-alone, 1.59±0.06, *n* = 25; SzSD: 1.92±0.04, *n* = 25; unpaired Welch’s t-test, \*\*\*\**P* < 0.0001. *Right:* Power (norm.): Two-way ANOVA show main effects of frequency band (*F* (2, 141) = 12.07; *P* < 0.0001) and event type (F (1, 141) = 15.66; *P* = 0.0001). Sz-alone vs SzSD: 1-10 Hz, *P* = 0.4131; 10-25 Hz, \*\**P* = 0.0014; 25-80 Hz, \*\**P* = 0.0065. For mean±SEM and multiple comparisons, see Supplementary Table 9. (**C**) SzSD recorded in ipsilateral somatosensory cortex (ipsi. SCtx) show higher coastline (C, *left*) and power (C, *right*) compared to seizure alone events. *Left:* Coastline (norm.): Sz-alone, 1.59±0.05, *n* = 35; SzSD: 1.89±0.04, *n* = 36; unpaired Welch’s t-test, \*\*\*\**P* < 0.0001. *Right:* Power (norm.): Two-way ANOVA show revealed a significant interaction between the factors (*F* (2, 204) = 5.653; *P* = 0.0041) and the main effects of frequency band (*F* (2, 204) = 11.08; p<0.0001) and event type (*F* (1, 204) = 22.42; *P* < 0.0001). Sz-alone vs SzSD: 1-10 Hz, *P* = 0.9312.; 10-25 Hz, \*\*\*\**P* < 0.0001; 25-80 Hz, \*\*\**P* = 0.0007. For mean±SEM and multiple comparisons, see Supplementary Table 10. (**D**) SzSD recorded in ipsilateral motor cortex (ipsi. MCtx) show higher coastline (D, *left*) and power (D, *right*) compared to seizure alone events. *Left:* Coastline (norm.): Sz-alone, 1.44±0.04, *n* = 35; SzSD: 1.67±0.03, *n* = 36; unpaired Mann-Whitney’s t-test, \*\*\*\**P* < 0.0001. *Right:* Power (norm.): Two-way ANOVA show main effects of frequency band (*F* (2, 204) = 15.26; *P* < 0.0001) and event type (*F* (1, 204) = 45.10; *P* < 0.0001). Sz-alone vs SzSD: 1-10 Hz, \*\**P* = 0.002; 10-25 Hz, \*\*\*\**P* < 0.0001; 25-80 Hz, \*\*\**P* = 0.0008. For mean±SEM and multiple comparisons, see Supplementary Table 11.

**Supplementary Figure 8.**
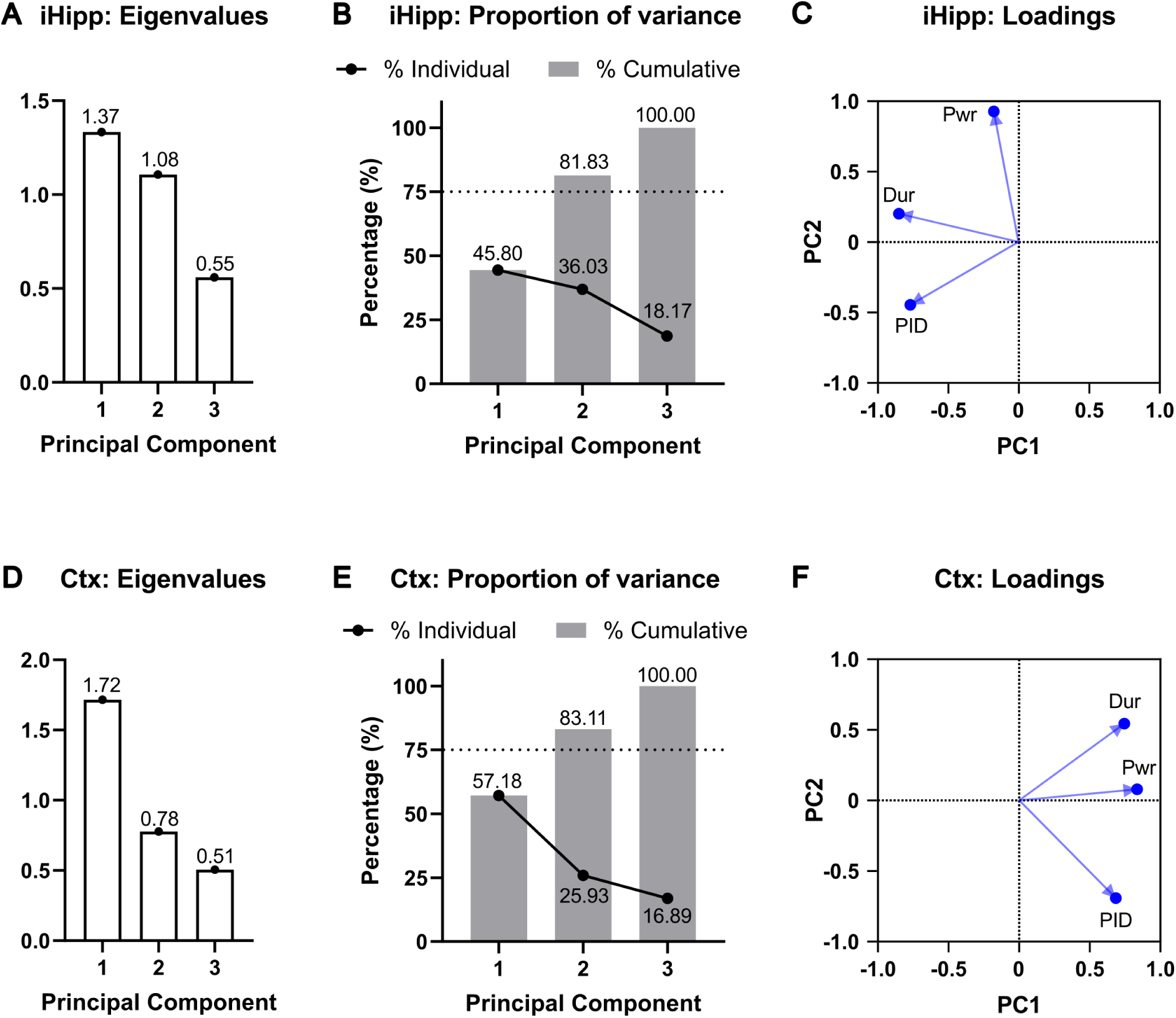
Principal component analysis of features of spontaneous seizure (power and duration) and postictal period (postictal depresssion) in hippocampal and cortical recordings in Kir4.1-cKO. (**A-C**) Principal component analysis (PCA) of ipsilateral hippocampal (iHipp) seizure events. (**A**) Eigenvalues for the first three principal components (PC). (**B**) Percentage variance explained by cumulative (gray bars) and individual PCs (black line). The first two PCs explained 81.83% of the total variance. Dotted line represnets 75% variance. (**C**) Loading plot showing the contribution of seizure power (Pwr), seizure duration (Dur) and postictal depression duration (PID) to PC1 and PC2. (**D-F**) PCA of cortical (Ctx) seizure events. (**D**) Eigenvalues for the first three PCs. (**E**) Percentage variance explained by by cumulative (gray bars) and individual PCs (black line). The first two PCs explained 83.11% of the total variance. Dotted line represnets 75% variance. (**F**) Loading plot showing the contribution of Pwr, Dur and PID to PC1 and PC2.

**Supplementary Figure 9.**
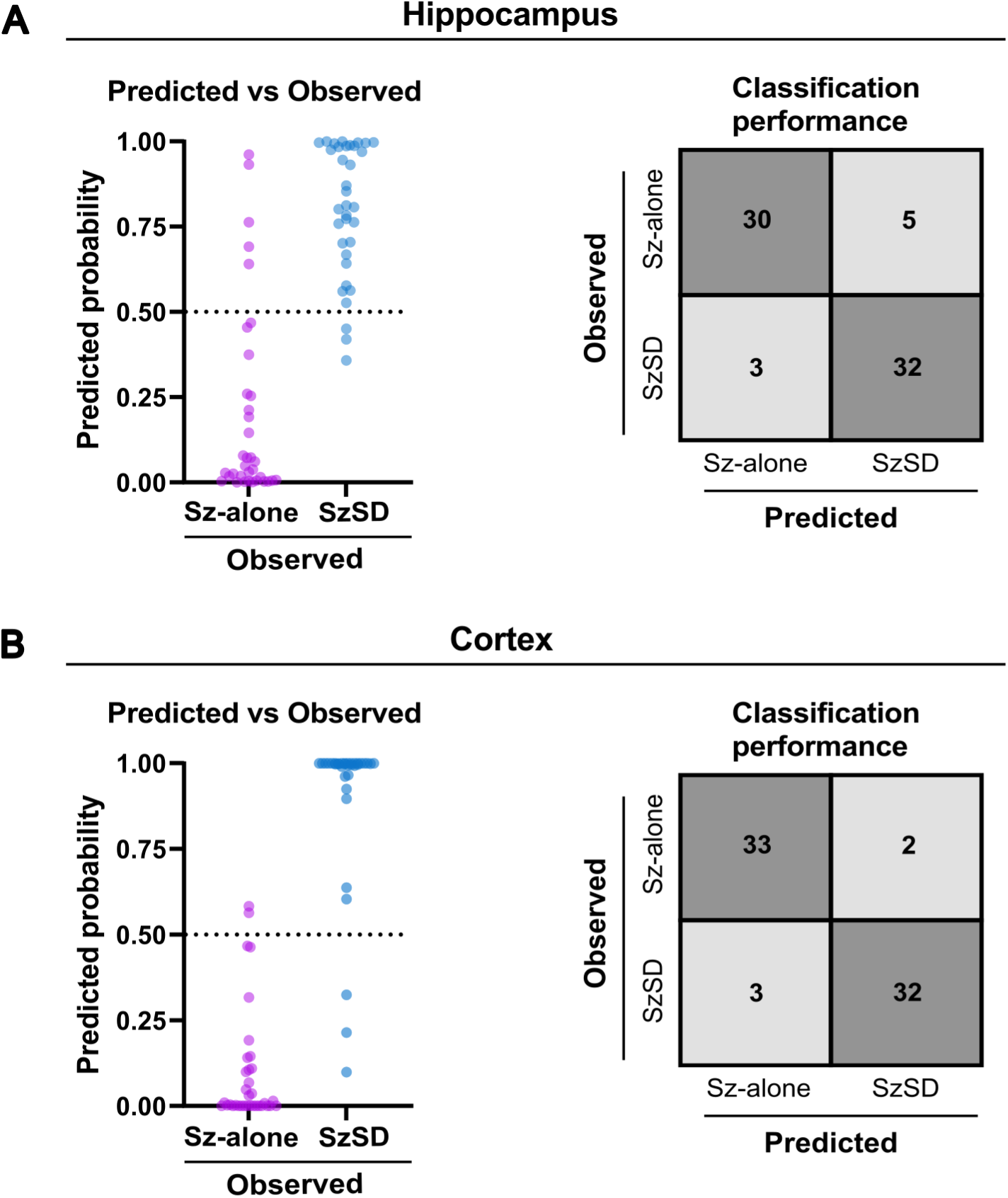
Logistic regression-based classification of SD-associated seizures using electrophysiological features obtainable from conventional AC-coupled recordings. (**A**) Hippocampal recordings. Left, predicted probabilities of classification as a seizure associated with spreading depolarization (SzSD) generated by the multivariable logistic regression model incorporating seizure power, seizure duration, and postictal depression. Each point represents a single seizure event. The dashed line indicates the classification threshold (0.5), with events above the threshold classified as SzSD and those below classified as seizure-alone (Sz-alone). Right, confusion matrix summarising model performance, with an overall classification accuracy of 88.57%. Correct classification rates were 85.71% for Sz-alone events (30/35) and 91.43% for SzSD events (32/35). (**B**) Cortical recordings. Left, predicted probabilities of classification as SzSD generated by the multivariable logistic regression model. Each point represents a single seizure event. The dashed line indicates the classification threshold (0.5). Right, confusion matrix summarising model performance, with an overall classification accuracy of 92.86%. Correct classification rates were 94.29% for Sz-alone events (33/35) and 91.43% for SzSD events (32/35).

**Supplementary Figure 10.**
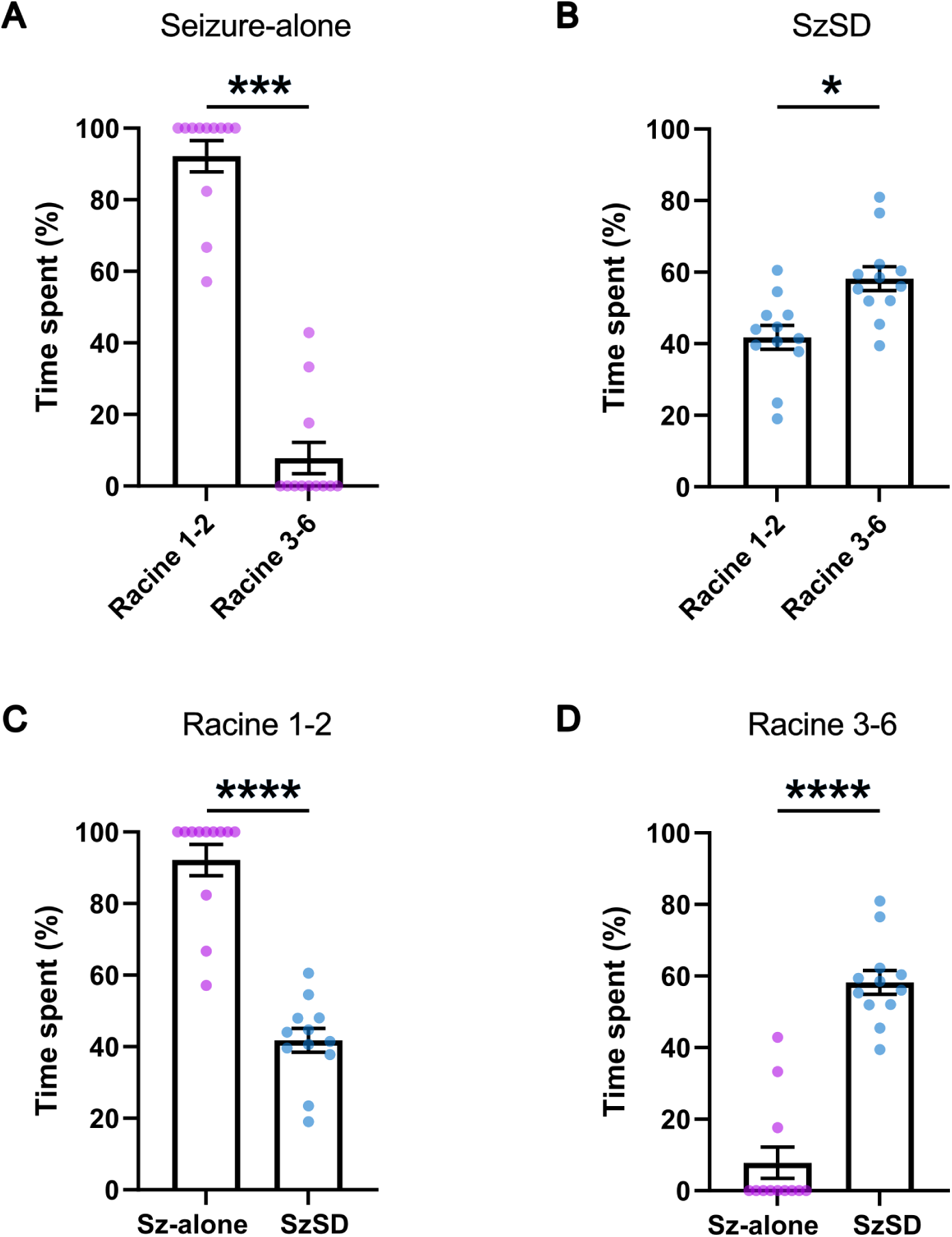
Seizures with spreading depolarizations show prolonged convulsive behavioral severity. (**A**) Seizure-alone (Sz-alone) events spent the majority of seizure duration in non-convulsive Racine stages (stages 1-2). Percentage of total seizure duration spent in Racine stages 1-2 and stages 3-6 was 92.18±4.37% and 7.82±4.38%, respectively (*n* = 12 events; Wilcoxon matched-pairs signed-rank test, \*\*\**P* = 0.0005). (**B**) Seizures associated with spreading depolarization (SzSD) spent a substantially greater proportion of seizure duration in convulsive Racine stages (stages 3-6). Percentage of total seizure duration spent in Racine stages 1-2 and stages 3-6 was 41.82±3.35% and 58.18±3.35%, respectively (*n* = 12 events; paired t-test, \**P* = 0.0326). (**C**) Sz-alone events spent a greater proportion of seizure duraion in non-conculsive Racine stages (stages1-2) compared with SzSD events (Sz, 92.18±4.37%; SzSD, 41.82±3.35%; *n* = 12 events/group; Mann-Whitney test, \*\*\*\**P* < 0.0001). (**D**) SzSD events spent a greater proportion of seizure duraion in the conculsive Racine stages (stages3-6) compared with seizure-alone events (Sz, 7.82±4.37%; SzSD, 58.1±3.35%; *n* = 12 events/group; Mann-Whitney test, \*\*\*\**P* < 0.0001).

**Supplementary Figure 11.**
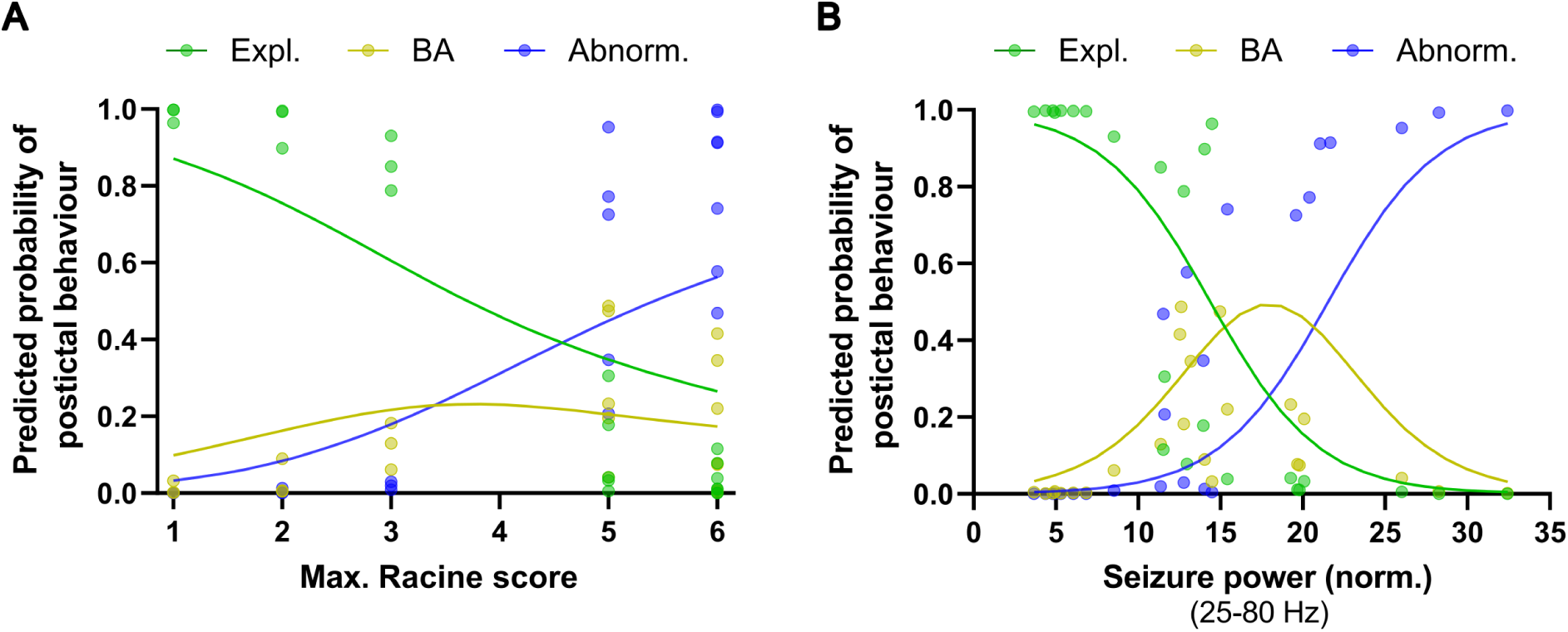
Seizure severity predicts the severity of postictal behavioural impairment. Ordinal logistic regression analysis showing the relationship between maximum seizure Racine score (**A**), seizure-associated spectral power (**B**), and postictal behavioural severity (exploration, behavioural arrest, and abnormal behaviours). Solid lines represent direct fitted probabilities derived from the ordinal logistic regression model, while circle markers indicate model-predicted probabilities at observed event values. Both maximum Racine score and seizure power were significant predictors of postictal behavioural category (overall model: χ²(2) = 29.2, *P* <0.001; *n* = 24 events). Increasing maximum Racine score was associated with greater odds of more severe postictal behavioural impairment (odds ratio (OR) = 3.46, 95% CI: 1.41-14.78, *P* = 0.029). Higher seizure power was also associated with increased odds of more severe postictal behavioural impairment (OR = 1.35, 95% CI: 1.02-2.15, *P* = 0.094). Likelihood ratio tests demonstrated significant effects of maximum Racine score (χ² = 8.38, *P* = 0.004) and seizure power (χ² = 4.51, *P* = 0.034).

**Supplementary Figure 12.**
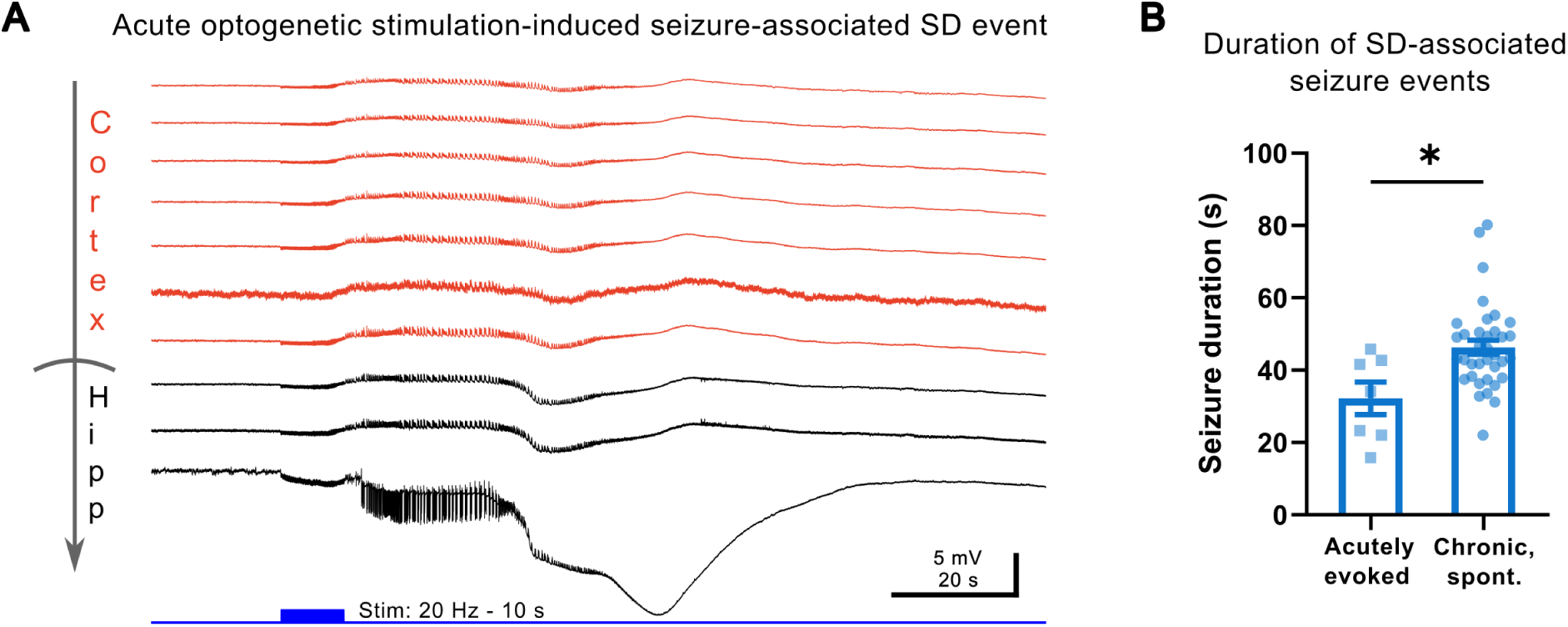
Duration of acutely evoked and chronic spontaneous SD-associated seizures. (**A**) Representative recordings showing optogenetically induced seizures associated with spreading depolarization (SD). SD emerged within the same hippocampal network that generated the seizure. Traces (top to bottom) represent recordings spanning from the superficial cortical layers (red) to the hippocampal region (black). (**B**) SD-associated seizures were shorter in duration during acutely evoked events compared with chronically recorded spontaneous seizures. Seizure duration: Evoked seizures, 32.20±4.48 s, *n* = 7; spontaneous seizures, 46.33±1.98 s, *n* = 36; Mann-Whitney test, \**P* = 0.0106.

**Supplementary Figure 13.**
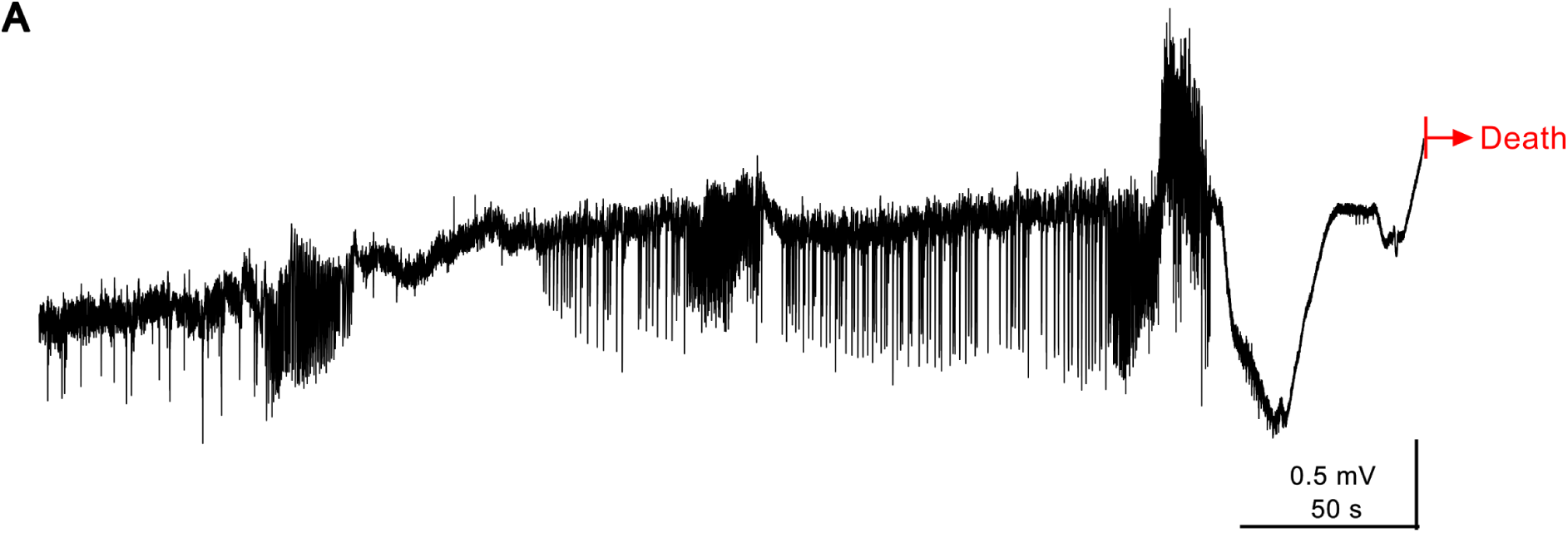
A terminal seizure cluster associated with spreading depolarization in a Kir4.1-cKO mouse. (**A**) Representative seizure cluster from a Kir4.1-cKO mouse in which the cluster-ending seizure was associated with spreading depolarization and followed by death. Recording was interrupted near the end of the trace (vertical red line) by animal facility staff to confirm death of the mouse.

## Supplementary movie legends

**Supplementary movie S1.**

Video clip time-locked to electrophysiological recordings showing a mouse during the final phase of a seizure-alone event and the subsequent postictal period. The mouse exhibits clonic seizure activity followed by rapid recovery and resumption of exploratory behaviour. All supplementary movies are from the same animal.

**Supplementary movie S2.**

Video clip time-locked to electrophysiological recordings showing an SD-associated seizure in the same animal as in Supplementary movie S1. The seizure progresses to Racine stage 6. Note, Supplementary movies S2-S5 show sequential stages of the same SD-associated seizure event and its postictal period.

**Supplementary movie S3.**

Video clip time-locked to electrophysiological recordings showing the late phase of the SD-associated seizure shown in Supplementary movie S2 and the transition into the postictal period. The clip captures seizure termination and the onset of postictal behaviours.

**Supplementary movie S4.**

Video clip time-locked to electrophysiological recordings showing the postictal period following the SD-associated seizure shown in Supplementary movies S2 and S3. The mouse remains behaviourally immobile approximately 60 minutes after seizure termination.

**Supplementary movie S5.**

Video clip time-locked to electrophysiological recordings showing a later stage of the postictal period following the SD-associated seizure shown in Supplementary movies S2-S4. The mouse exhibits movements and grooming behaviour, indicators of behavioural recovery from postictal behavioural arrest.

