## Supplementary Tables for "Focal astrocyte Kir4.1 loss drives seizures, spreading depolarizations and postictal impairments"

**Supplementary Table 1: Electrical stimulation-induced DC shift (mV) *ex vivo***

| DC shift (mV) | | | |
|---|---|---|---|
| Mean±SEM | 5 Hz (*n*) | 20 Hz (*n*) | 50 Hz (*n*) |
| Control | 0.15±0.02 (5) | 0.21±0.03 (5) | 0.28±0.03 (5) |
| cKO | 0.18±0.01 (6) | 0.38±0.05 (6) | 0.77±0.10 (6) |
| **Two-way ANOVA** | | | |
| | *F* (DFn, DFd) | *P* value | |
| Interaction | *F* (2, 27) = 9.823 | 0.0006 | |
| Stim. Frequency | *F* (2, 27) = 22.44 | <0.0001 | |
| Treatment groups | *F* (1, 27) = 27.28 | <0.0001 | |
| **Tukey's Multiple comparison test** | | | |
| **Adjusted *P*-value** | 5 Hz | 20 Hz | 50 Hz |
| Control vs cKO | 0.7394 | 0.0329 | <0.0001 |
| **Adjusted *P*-value** | 5 Hz vs 20 Hz | 5 Hz vs 50 Hz | 20 Hz vs 50 Hz |
| Control | 0.7808 | 0.2885 | 0.6652 |
| cKO | 0.0279 | <0.0001 | <0.0001 |

cKO = Kir4.1-conditional knockout
*n* values represent number of slices

**Supplementary Table 2: GINKO responses *ex vivo* - Maximum amplitude (dF/F0)**

| GINKO amplitude (dF/F0) | | | |
|---|---|---|---|
| Mean±SEM | 5 Hz (*n*) | 20 Hz (*n*) | 50 Hz (*n*) |
| Control | 4.15±0.27 (5) | 11.33±1.38 (5) | 12.96±1.03 (5) |
| cKO | 9.74±2.51 (5) | 21.78±3.74 (5) | 27.60±3.40 (5) |
| **Two-way ANOVA** | | | |
| | *F* (DFn, DFd) | *P* value | |
| Interaction | *F* (2, 24) = 1.763 | 0.1931 | |
| Stim. Frequency | *F* (2, 24) = 16.31 | <0.0001 | |
| Treatment groups | *F* (1, 24) = 27.02 | <0.0001 | |
| **Tukey's Multiple comparison test** | | | |
| **Adjusted *P*-value** | 5 Hz | 20 Hz | 50 Hz |
| Control vs cKO | 0.1137 | 0.0053 | 0.0002 |
| **Adjusted *P*-value** | 5 Hz vs 20 Hz | 5 Hz vs 50 Hz | 20 Hz vs 50 Hz |
| Control | 0.1096 | 0.0413 | 0.8815 |
| cKO | 0.0047 | <0.0001 | 0.2235 |

cKO = Kir4.1 conditional knockout
*n* values represent number of slices

**Supplementary Table 3: Transduction volume ($mm^3$) of Ast-Cre-mCherry viral vector**

| **Transduction volume ($mm^3$)** | | | | |
|---|---|---|---|---|
| | Left hippocampus (L.Hipp) | Left cortex (L.Ctx) | Right hippocampus (R.Hipp)[$] | Right cortex (R.Ctx) |
| Mean±SEM | 0.22±0.11 | 0.05±0.11 | 3.09±0.19 | 0.92±0.32 |
| *n* | 7 | 7 | 7 | 7 |
| **Two-way RM ANOVA** | | | | |
| | *F* (DFn, DFd) | | *P* value | |
| Interaction | *F* (1, 6) = 43.14 | | 0.0006 | |
| Region | *F* (1, 6) = 184.5 | | <0.0001 | |
| Hemisphere | *F* (1, 6) = 80.86 | | <0.0001 | |
| **Tukey's Multiple comparison test** | | | | |
| | **Adjusted *P*-value** | | | |
| R.Hipp vs L.Hipp | <0.0001 | | | |
| R.Hipp vs R.Ctx | 0.0002 | | | |
| R.Hipp vs L.Ctx | <0.0001 | | | |
| L.Hipp vs R.Ctx | 0.0626 | | | |
| L.Hipp vs L.Ctx | 0.8556 | | | |
| R.Ctx vs L.Ctx | 0.0256 | | | |

cKO = Kir4.1 conditional knockout.
[$]Ast-Cre-mCherry was injected into the right hippocampus.
*n* values represent number of mice

**Supplementary Table 4: Optogenetic stimulation-induced DC shift (mV) *in vivo***

| **DC shift (mV)** | | | |
|---|---|---|---|
| Mean±SEM | 5 Hz (*n*) | 20 Hz (*n*) | 50 Hz (*n*) |
| Control | 0.12±0.05 (4) | 0.23±0.11 (4) | 0.54±0.22 (4) |
| cKO | 0.58±0.13 (5) | 1.79±0.53 (5) | 2.98±0.66 (5) |
| **Two-way ANOVA** | | | |
| | *F* (DFn, DFd) | | *P* value |
| Interaction | *F* (2, 21) = 3.008 | | 0.0710 |
| Stim. Frequency | *F* (2, 21) = 5.996 | | 0.0087 |
| Treatment groups | *F* (1, 21) = 20.06 | | 0.0002 |
| **Tukey's Multiple comparison test** | | | |
| **Adjusted *P*-value** | 5 Hz | 20 Hz | 50 Hz |
| Control vs cKO | 0.4376 | 0.0130 | 0.0004 |
| **Adjusted *P*-value** | 5 Hz vs 20 Hz | 5 Hz vs 50 Hz | 20 Hz vs 50 Hz |
| Control | 0.9821 | 0.7777 | 0.8727 |
| cKO | 0.0871 | 0.0007 | 0.0964 |

cKO = Kir4.1 conditional knockout
*n* values represent number of trials

**Supplementary Table 5: Postictal activity recovery (percentage power change) after optogenetic stimulation-induced seizures and SzSD in Kir4.1-cKO *in vivo*.**

| Percentage power change (Δ%) | | | | | | | | |
|---|---|---|---|---|---|---|---|---|
| Mean±SEM | Bfr | Sz | r1 | r2 | r3 | r4 | r5 | r6 |
| Seizures (*n*=4) | 0.00±0.00 | 444.12±109.57 | 51.67±30.96 | -7.59±10.35 | 5.54±5.60 | 1.02±5.44 | 7.75±5.66 | 13.03±9.43 |
| SzSD (*n*=7) | 0.00±0.00 | 738.58±258.34 | -54.16±14.36 | -42.64±9.81 | -33.13±9.04 | -23.45±9.45 | -20.31±7.90 | -15.15±6.96 |
| **Two-way RM ANOVA** | | | | | | | | |
| | *F* (DFn, DFd) | | | | *P* value | | | |
| Interaction | *F* (1.013, 9.115) = 0.8912 | | | | 0.3709 | | | |
| Recovery time | *F* (1.013, 9.115) = 10.95 | | | | 0.0088 | | | |
| Event type | *F* (1, 9) = 0.009 | | | | 0.9241 | | | |
| **Tukey's Multiple comparison test** | | | | | | | | |
| **Adjusted *P*-value** | Bfr | Sz | r1 | r2 | r3 | r4 | r5 | r6 |
| Sz vs SzSD | - | 0.3253 | 0.0325 | 0.0406 | 0.0056 | 0.0524 | 0.0180 | 0.0511 |

'Bfr', Before seizure
'Sz', Seizure-alone events
'r' in r1-6, recovery
'SzSD', SD-associated seizure events
*n* values represent number of trials

**Supplementary Table 6: Postictal activity recovery (percentage power change) after spontaneous seizures and SzSD in Kir4.1-cKO *in vivo*.**

| Percentage power change (Δ%) | | | | | | | | |
|---|---|---|---|---|---|---|---|---|
| Mean±SEM | Bfr (*n*) | Sz (*n*) | r1 (*n*) | r2 (*n*) | r3 (*n*) | r4 (*n*) | r5 (*n*) | r6 (*n*) |
| Seizures | 0.00±0.00 (35) | 623.10±41.10 (35) | -4.91±10.75 (35) | -7.30±9.05 (32) | 15.60±14.75 (30) | 31.38±33.80 (26) | 30.24±22.42 (26) | 22.97±19.50 (24) |
| SzSD | 0.00±0.00 (35) | 710.37±73.73 (35) | -59.85±3.95 (35) | -66.98±3.27 (35) | -40.85±7.42 (35) | -23.09±6.07 (35) | -22.41±5.78 (35) | -15.91±5.62 (34) |
| **Linear Mixed Effects using restricted maximum likelihood (REML)** | | | | | | | | |
| | *F* (DFn, DFd) | | | | *P* value | | | |
| Interaction | *F* (1.309, 81.90) = 2.284 | | | | 0.127 | | | |
| Recovery time | *F* (1.309, 81.90) = 215.4 | | | | <0.0001 | | | |
| Event type | *F* (1, 68) = 2.713 | | | | 0.1041 | | | |
| **Tukey's Multiple comparison test** | | | | | | | | |
| **Adjusted *P*-value** | Bfr | Sz | r1 | r2 | r3 | r4 | r5 | r6 |
| Sz vs SzSD | - | 0.3059 | <0.0001 | <0.0001 | 0.0014 | 0.1245 | 0.0307 | 0.0661 |

'Bfr', Before seizure
'Sz', Seizure-alone events
'r' in r1-6, recovery
'SzSD', SD-associated seizure events
*n* values represent number of events

**Supplementary Table 7: Quantification of seizure duration – Cortex vs Hippocampus**

| Seizure duration (s) | | | | |
|---|---|---|---|---|
| Region | iHipp | | Ctx | |
| | Sz (*n*) | SzSD (*n*) | Sz (*n*) | SzSD (*n*) |
| Mean±SEM | 32.68±1.57 (35) | 46.33±1.98 (36) | 34.06±1.53 (35) | 46.61±2.02 (36) |
| **Two-way ANOVA** | | | | |
| | *F* (DFn, DFd) | | *P* value | |
| Interaction | *F* (1, 138) = 0.09351 | | 0.7602 | |
| Frequency bands | *F* (1, 138) = 53.21 | | <0.0001 | |
| Event type | *F* (1, 138) = 0.2133 | | 0.6449 | |
| **Tukey's Multiple comparison test** | | | | |
| | | | **Adjusted *P*-value** | |
| Sz: iHipp vs. Sz: Ctx | | | 0.9493 | |
| Sz: iHipp vs. SzSD: iHipp | | | <0.0001 | |
| Sz: iHipp vs. SzSD: Ctx | | | <0.0001 | |
| Sz: Ctx vs. SzSD: iHipp | | | <0.0001 | |
| Sz: Ctx vs. SzSD: Ctx | | | <0.0001 | |
| SzSD: iHipp vs. SzSD: Ctx | | | 0.9995 | |

'Sz', Seizure-alone events
'SzSD', SD-associated seizure events
'iHipp', ipsilateral hippocampus
'Ctx', cortex
*n* values represent number of events

**Supplementary Table 8: Quantification of seizure power - ipsilateral hippocampus**

| Seizure power (norm.) | | | |
|---|---|---|---|
| Mean±SEM | 1-10 Hz | 10-25 Hz | 25-80 Hz |
| Seizure-alone (*n*=35) | 19.20±1.98 | 14.70±2.31 | 10.63±1.49 |
| SzSD (*n*=36) | 27.66±5.99 | 20.41±2.75 | 13.43±1.56 |
| **Two-way ANOVA** | | | |
| | *F* (DFn, DFd) | *P* value | |
| Interaction | *F* (2, 207) = 0.4129 | 0.6623 | |
| Frequency bands | *F* (2, 207) = 6.68 | 0.0015 | |
| Event type | *F* (1, 207) = 4.94 | 0.0274 | |
| **Tukey's Multiple comparison test** | | | |
| **Adjusted *P*-value** | 1-10 Hz | 10-25 Hz | 25-80 Hz |
| Seizure-alone vs SzSD | 0.0563 | 0.1967 | 0.5266 |
| **Adjusted *P*-value** | 1-10 Hz vs 10-25 Hz | 1-10 Hz vs 25-80 Hz | 10-25 Hz vs 25-80 Hz |
| Seizure-alone | 0.5692 | 0.1334 | 0.6317 |
| SzSD | 0.2246 | 0.0038 | 0.2509 |

'SzSD', SD-associated seizure events
*n* values represent number of events

**Supplementary Table 9: Quantification of seizure power - contralateral somatosensory cortex**

| Seizure power (norm.) | | | |
|---|---|---|---|
| Mean±SEM | 1-10 Hz | 10-25 Hz | 25-80 Hz |
| Seizure-alone (*n*=25) | 24.87±2.50 | 19.11±2.77 | 9.51±1.07 |
| SzSD (*n*=25) | 27.89±3.82 | 29.98±3.08 | 19.13±1.19 |
| **Two-way ANOVA** | | | |
| | F (DFn, DFd) | *P* value | |
| Interaction | *F* (2, 144) = 1.32 | 0.2710 | |
| Frequency bands | *F* (2, 144) = 12.51 | <0.0001 | |
| Event type | *F* (1, 144) = 13.64 | 0.0003 | |
| **Tukey's Multiple comparison test** | | | |
| **Adjusted *P*-value** | 1-10 Hz | 10-25 Hz | 25-80 Hz |
| Seizure-alone vs SzSD | 0.4126 | 0.0036 | 0.0098 |
| **Adjusted *P*-value** | 1-10 Hz vs 10-25 Hz | 1-10 Hz vs 25-80 Hz | 10-25 Hz vs 25-80 Hz |
| Seizure-alone | 0.2631 | 0.0001 | 0.0268 |
| SzSD | 0.8367 | 0.0481 | 0.0102 |

'SzSD', SD-associated seizure events
*n* values represent number of events

**Supplementary Table 10: Quantification of seizure power - ipsilateral somatosensory cortex**

| Seizure power (norm.) | | | |
|---|---|---|---|
| Mean±SEM | 1-10 Hz | 10-25 Hz | 25-80 Hz |
| Seizure-alone (*n*=35) | 21.70±2.14 | 13.72±1.73 | 9.89±1.36 |
| SzSD (*n*=36) | 21.51±1.85 | 24.26±2.20 | 16.96±1.57 |
| **Two-way ANOVA** | | | |
| | *F* (DFn, DFd) | *P* value | |
| Interaction | *F* (2, 207) = 4.44 | 0.0129 | |
| Frequency bands | *F* (2, 207) = 10.36 | <0.0001 | |
| Event type | *F* (1, 207) = 15.00 | 0.0001 | |
| **Tukey's Multiple comparison test** | | | |
| **Adjusted *P*-value** | 1-10 Hz | 10-25 Hz | 25-80 Hz |
| Seizure-alone vs SzSD | 0.9424 | <0.0001 | 0.007 |
| **Adjusted *P*-value** | 1-10 Hz vs 10-25 Hz | 1-10 Hz vs 25-80 Hz | 10-25 Hz vs 25-80 Hz |
| Seizure-alone | 0.0072 | <0.0001 | 0.3094 |
| SzSD | 0.5370 | 0.1835 | 0.0141 |

'SzSD', SD-associated seizure events
*n* values represent number of events

**Supplementary Table 11: Quantification of seizure power - ipsilateral motor cortex**

| Seizure power (norm.) | | | |
|---|---|---|---|
| Mean±SEM | 1-10 Hz | 10-25 Hz | 25-80 Hz |
| Seizure-alone (*n*=35) | 10.44±1.09 | 9.73±1.23 | 4.82±0.48 |
| SzSD (*n*=36) | 15.41±1.34 | 18.14±1.78 | 10.38±0.85 |
| **Two-way ANOVA** | | | |
| | *F* (DFn, DFd) | P value | |
| Interaction | *F* (2, 207) = 1.17 | 0.3128 | |
| Frequency bands | *F* (2, 207) = 15.95 | <0.0001 | |
| Event type | *F* (1, 207) = 45.20 | <0.0001 | |
| **Tukey's Multiple comparison test** | | | |
| **Adjusted *P*-value** | 1-10 Hz | 10-25 Hz | 25-80 Hz |
| Seizure-alone vs SzSD | 0.0039 | <0.0001 | 0.0013 |
| **Adjusted *P*-value** | 1-10 Hz vs 10-25 Hz | 1-10 Hz vs 25-80 Hz | 10-25 Hz vs 25-80 Hz |
| Seizure-alone | 0.9090 | 0.0035 | 0.0130 |
| SzSD | 0.2419 | 0.0093 | <0.0001 |

'SzSD', SD-associated seizure events
*n* values represent number of events

**Supplementary Table 12: Quantification of seizure power – Cortex vs Hippocampus**

| Seizure power (norm.) \| 25-80 Hz | | | | |
|---|---|---|---|---|
| Region | iHipp | | Ctx | |
| | Sz (*n*) | SzSD (*n*) | Sz (*n*) | SzSD (*n*) |
| Mean±SEM | 13.56±1.48 (35) | 14.45±1.53 (36) | 9.20±0.85 (35) | 16.35±1.05 (36) |
| **Two-way ANOVA** | | | | |
| | *F* (DFn, DFd) | | *P* value | |
| Interaction | *F* (1, 138) = 6.143 | | 0.0144 | |
| Event type | *F* (1, 138) = 10.15 | | 0.0018 | |
| Region | *F* (1, 138) = 0.9521 | | 0.3309 | |
| **Tukey's Multiple comparison test** | | | | |
| | | **Adjusted *P*-value** | | |
| Sz: iHipp vs. Sz: Ctx | | 0.0770 | | |
| Sz: iHipp vs. SzSD: iHipp | | 0.9590 | | |
| Sz: iHipp vs. SzSD: Ctx | | 0.4035 | | |
| Sz: Ctx vs. SzSD: iHipp | | 0.0198 | | |
| Sz: Ctx vs. SzSD: Ctx | | 0.0006 | | |
| SzSD: iHipp vs. SzSD: Ctx | | 0.7082 | | |

'Sz', Seizure-alone events
'SzSD', SD-associated seizure events
'iHipp', ipsilateral hippocampus
'Ctx', cortex
*n* values represent number of events

**Supplementary Table 13: Quantification of Postictal depression – Cortex vs Hippocampus**

| **Postictal depression (1-40 Hz power ratio: pre-Sz/post-Sz)** | | | | |
|---|---|---|---|---|
| Region | iHipp | | Ctx | |
| | Sz (*n*) | SzSD (*n*) | Sz (*n*) | SzSD (*n*) |
| Mean±SEM | 1.33±0.13 (35) | 3.56±0.41 (35) | 1.22±0.10 (35) | 3.27±0.40 (35) |
| **Two-way ANOVA** | | | | |
| | *F* (DFn, DFd) | | *P* value | |
| Interaction | *F* (1, 136) = 0.0990 | | 0.7535 | |
| Event type | *F* (1, 136) = 51.64 | | <0.0001 | |
| Region | *F* (1, 136) = 0.4721 | | 0.4905 | |
| **Tukey's Multiple comparison test** | | | | |
| | | **Adjusted *P*-value** | | |
| Sz: iHipp vs. Sz: Ctx | | 0.9934 | | |
| Sz: iHipp vs. SzSD: iHipp | | <0.0001 | | |
| Sz: iHipp vs. SzSD: Ctx | | <0.0001 | | |
| Sz: Ctx vs. SzSD: iHipp | | <0.0001 | | |
| Sz: Ctx vs. SzSD: Ctx | | <0.0001 | | |
| SzSD: iHipp vs. SzSD: Ctx | | 0.4905 | | |

'Sz', Seizure-alone events
'SzSD', SD-associated seizure events
'iHipp', ipsilateral hippocampus
'Ctx', cortex
*n* values represent number of events

**Supplementary Table 14: Correlation Matrix: ipsilateral hippocampus**

| **Spearman r correlation Matrix** (*n* =70) | | | |
|---|---|---|---|
| **Correlation values** | Power | Duration | PID |
| Power | 1.00 | 0.14 | -0.10 |
| Duration | 0.14 | 1.00 | 0.40 |
| PID | -0.10 | 0.40 | 1.00 |
| ***P*-values** | | | |
| ***P*-values** | Power | Duration | PID |
| Power | | 0.2386 | 0.4316 |
| Duration | 0.2386 | | 0.0005 |
| PID | 0.4316 | 0.0005 | |

'PID', Postictal depression
*n* values represent number of events

**Supplementary Table 15: Correlation Matrix: Cortex**

| **Spearman r correlation Matrix** (*n* =70) | | | |
|---|---|---|---|
| **Correlation values** | Power | Duration | PID |
| Power | 1.00 | 0.46 | 0.39 |
| Duration | 0.46 | 1.00 | 0.34 |
| PID | 0.39 | 0.34 | 1.00 |
| ***P*-values** | | | |
| ***P*-values** | Power | Duration | PID |
| Power | | <0.0001 | 0.0007 |
| Duration | <0.0001 | | 0.0044 |
| PID | 0.0007 | 0.0044 | |

'PID', Postictal depression
*n* values represent number of events

**Supplementary Table 16: Odd ratios and 95% Confidence intervals**

| **Odd ratios and 95% Confidence intervals** | | | | |
|---|---|---|---|---|
| | **iHipp** (*n* =70) | | **Ctx** (*n* =70) | |
| | Odds ratio, [95% CI] | *P* value | Odds ratio, [95% CI] | *P* value |
| PID | $1.48 \times 10^3$, [$3.88 \times 10^1$, $3.45 \times 10^5$] | < 0.0001 | $5.83 \times 10^8$, [$5.44 \times 10^{-18}$, $5.84 \times 10^{-5}$] | < 0.0001 |
| Dur | $8.30 \times 10^5$, [$6.68 \times 10^2$, $9.48 \times 10^9$] | < 0.0001 | $1.31 \times 10^8$, [$3.45 \times 10^2$, $6.61 \times 10^{16}$] | 0.0017 |
| Pwr | 5.43, [$2.8 \times 10^{-2}$, $1.90 \times 10^2$] | 0.2751 | $1.05 \times 10^4$, [$8.14 \times 10^1$, $6.41 \times 10^8$] | 0.0074 |

'PID', Postictal depression
'Dur', Seizure duration
'Pwr', Seizure power
'iHipp', ipsilateral hippocampus
'Ctx', cortex
*n* values represent number of events

**Supplementary Table 17: Time spent (%) in each Racine stage during spontaneous seizures (seizure alone and SzSD) in Kir4.1-cKO.**

| **Time (%)** | | | | | | |
|---|---|---|---|---|---|---|
| Mean±SEM | Racine 1 | Racine 2 | Racine 3 | Racine 4 | Racine 5 | Racine 6 |
| Sz-alone (*n*=12) | 78.48±8.98 | 13.70±6.08 | 7.82±4.37 | 0.00±0.00 | 0.00±0.00 | 0.00±0.00 |
| SzSD (*n*=12) | 19.09±4.18 | 22.72±4.04 | 18.03±3.64 | 9.78±4.22 | 24.05±4.30 | 6.33±2.12 |
| **Two-way RM ANOVA** | | | | | | |
| | *F* (DFn, DFd) | | | *P* value | | |
| Interaction | $F$ (2.042, 44.92) = 19.71 | | | <0.0001 | | |
| Racine stage | $F$ (2.042, 44.92) = 24.77 | | | <0.0001 | | |
| Repeats | $F$ (1, 22) = 5.66 | | | 0.0265 | | |
| **Tukey's Multiple comparison test** | | | | | | |
| **Adjusted *P*-value** | Racine 1 | Racine 2 | Racine 3 | Racine 4 | Racine 5 | Racine 6 |
| Sz-alone vs SzSD | <0.0001 | 0.2314 | 0.0868 | 0.0407 | 0.0002 | 0.0123 |

'Sz-alone', Seizure-alone events
'SzSD', SD-associated seizure events
*n* values represent number of events

**Supplementary Table 18: Quantification of Postictal behaviour (percentage of total time) during the 120-second window after the end of the electrographic seizure**

| Time (%) | | | |
|---|---|---|---|
| Mean±SEM | Exploring | Behavioural arrest | Abnormal |
| Seizures-alone (*n*=12) | 98.33±0.95 | 1.18±0.68 | 0.49±0.49 |
| SzSD (*n*=12) | 12.64±4.61 | 28.96±5.66 | 58.40±6.07 |
| **Two-way ANOVA** | | | |
| | *F* (DFn, DFd) | P value | |
| Interaction | $F$ (2, 66) = 187.4 | <0.0001 | |
| Behaviour | $F$ (2, 66) = 54.87 | <0.0001 | |
| Event type | $F$ (1, 66) = 7.566e-009 | >0.9999 | |
| **Tukey's Multiple comparison test** | | | |
| **Adjusted *P*-value** | Exploring | Behavioural arrest | Abnormal |
| Seizures-alone vs SzSD | <0.0001 | <0.0001 | <0.0001 |
| **Adjusted p-value** | Behavioural arrest vs Abnormal | Behavioural arrest vs Exploring | Abnormal vs Exploring |
| Seizures-alone | 0.9914 | <0.0001 | <0.0001 |
| SzSD | <0.0001 | 0.012 | <0.0001 |

'SzSD', SD-associated seizure events
*n* values represent number of events
